# Evaluating the use of non-linear models in data-driven rescoring of peptide-spectrum matches

**DOI:** 10.64898/2026.07.10.737772

**Authors:** Alireza Nameni, Arthur Declercq, Ralf Gabriels, Sven Degroeve, Lennart Martens, Robbin Bouwmeester

**Affiliations:** CompOmics, VIB Center for Medical Biotechnology, VIB, Ghent 9052, Belgium; Department of Biomolecular Medicine, Faculty of Medicine and Health Sciences, Ghent University, Ghent 9052, Belgium. Department of Biomolecular Medicine, Ghent University, Ghent, Belgium; BioOrganic Mass Spectrometry Laboratory (LSMBO), IPHC UMR 7178, University of Strasbourg, CNRS, Strasbourg, 67000, France; Infrastructure Nationale de Protéomique ProFI − FR2048, Strasbourg, 67087, France

## Abstract

In mass spectrometry (MS)-based proteomics, computational tools match acquired tandem MS spectra to peptides from a sequence database. Machine learning increasingly supports this task through peptide-spectrum match (PSM) rescoring, in which a classifier, typically a linear semi-supervised model, refines the initial matching score. However, Mokapot allows the user to choose among different machine learning algorithms of increasing complexity, from the default linear support vector machine (LSVM) to random forest and XGBoost. Here, we use an entrapment approach to assess the effect of this increasing complexity on PSM identification and the accuracy of the estimated false discovery rate (FDR). We show that, while more complex models increase the number of identified PSMs at a fixed FDR threshold, this gain reflects a bias towards random matches from the target proteome database rather than genuine identifications. Indeed, for the most complex model, the entrapment FDR reaches 6.3% instead of the estimated 1% decoy FDR. This bias thus yields overly optimistic FDR estimates, indicating that model complexity in PSM rescoring must be carefully balanced against this overfitting risk.

## 1 Introduction

Liquid chromatography coupled to mass spectrometry (LC-MS) is the technology of choice for high throughput proteomics analysis^1^. LC-MS generates large volumes of raw signal data that require specialized computational tools for processing and interpretation. These tools match peptides from a target proteome database to acquired MS spectra (PSMs). These peptide-spectrum matches are evaluated by an assigned PSM score, representing the likelihood that the given peptide has generated the observed spectrum. A null distribution of PSM scores is then computed from decoy peptide sequences, which are generated by shuffling or reversing the protein or peptide sequences in the target proteome database. By matching these decoy sequences to spectra along with target sequences, statistical confidence estimates can be assigned to the obtained target PSMs^2^.

To increase identification rate of spectra acquired in data dependent acquisition (DDA) mode, and thus improve identification sensitivity, it has become common practice to post-process PSMs with machine learning. In this method, called rescoring, the model learns to optimally combine individual scoring metrics into a single compounded PSM score. The most common rescoring approach involves semi-supervised learning, where the model is trained using a small set of labeled examples alongside a larger pool of unlabeled data. In this approach, high-confidence target PSMs serve as initial positive training examples and decoy PSMs as initial negatives, while the remaining PSMs are iteratively classified and used to refine the model^3^. The final goal is to differentiate true target PSMs from false target PSMs with a classification model. PSMs are represented as feature vectors that, in addition to the PSM scores obtained from a search engine, include PSM features like MS mass errors (MS1 and MS2), number of matched peaks (MS1 and MS2), and peptide characteristics such as sequence length and charge state. Several tools implement this rescoring strategy^4^, most notably Percolator^3^ and Mokapot^5^. More recently, data-driven rescoring tools such as MS^2^Rescore^6^, MSBooster^7^, and Oktoberfest^8^ have extended this strategy by incorporating additional features derived from machine learning-based peptide property predictors, such as predicted fragment ion intensities and retention times, yielding further improvements in PSM identification rates^9^. Moreover, similar methods have also been adopted in several data independent acquisition (DIA) search algorithms^10^.

Mokapot’s rescoring implementation is particularly interesting as it allows for the selection of various machine learning algorithms with different degrees of model complexity^5^. In particular, it was shown that more complex models, such as the gradient tree boosting (XGBoost) algorithm^11^, increase the PSM identification rate. However, applying more complex models for PSM rescoring could also result in overfitting the target proteome database. In such cases, false target matches may have a higher probability of being assigned a positive PSM score than true decoy matches. This indicates that the model is overfitted to classify target peptides as positives, regardless of being true or false, which can result in overly optimistic FDR estimates. This concern may be further amplified in data-driven rescoring, where a large number of additional features can be incorporated into the rescoring model. Moreover, as these features are themselves derived from machine learning models, they could introduce additional biases against decoy peptides if these models were not trained correctly.

In this study, we investigate the potential for overfitting in rescoring when using models with increasing complexity. To do so, we adopted the entrapment sequence database approach from The *et al*.^*12*^ to compute the entrapment FDR. This procedure provides a means to assess any potential overfitting of the rescoring method and allows us to gain valuable insight into the relation between the degree of complexity of the model and the tendency to overfit the target proteome database.

## 2 MATERIALS AND METHODS

### 2.1 Spectrum files

We selected subsets of spectrum files from eleven publicly available LC-MS experiments to evaluate the different models: CD8 T cells (PXD000561, LTQ Orbitrap Velos Elite)^13^, Human heart tissue (PXD006675, Q Exactive)^14^, HEK293 cells (PXD001468, Q Exactive)^15^, Mouse brain tissue (PXD001250, Q Exactive)^16^, HeLa cells (PXD000612, Q Exactive)^17^, Mouse NIH/3T3 fibroblast cells (PXD004948, Q Exactive)^18^, gills and caeca from *Gammarus fossarum* (PXD040344, Q Exactive)^19^, *Solanum lycopersicum* (PXD004947, Q Exactive)^20^, *Bacillus subtilis* (PXD004565, Q Exactive)^21^, *Methanosarcina mazei* (PXD004325, Q Exactive)^22^, and *Saccharomyces cerevisiae* (PXD009815, Q Exactive)^23^. Raw data files were downloaded from PRIDE and are listed in Supplementary Table 1. These datasets were selected to represent a diverse range of organisms, sample types, and experimental conditions, thus providing a robust basis for evaluating the models.

### 2.2 Search engine settings

The spectrum files were searched against the corresponding organism’s protein sequence database (Supplementary Table 1), entrapment sequences, and contaminants using SearchGUI^24^(v4.3.1) with default settings, incorporating Andromeda^25^, Comet^26^, MS-GF+^27^, and MS Amanda^28^. *In silico* digestion settings were configured for trypsin without proline suppression, allowing for up to two missed cleavages, minimum peptide length of eight amino acids, and maximum length of 30 amino acids. Oxidation (M) and acetylation (protein N-term) were set as variable modifications, and carbamidomethyl (C) was set as fixed modification. The decoy strategy was set to reverse protein sequences and generated by SearchGUI, with a disabled FDR filter (set at 100%) to allow for downstream rescoring. The SearchGUI settings file is available on Zenodo.

### 2.3 Data processing

The identification results from different search engines were then converted to the open mzIdentML (.mzid) format using PeptideShaker (v3.0.1). Subsequently, MS^2^Rescore (v3.0.0b5)^6^ was used to add a comprehensive set of features including basic PSM attributes and search engine-generated scores, as well as features derived from predicted fragment ion intensities (MS^2^PIP v4.0.0) and retention times (DeepLC v2.2.22). The results were then exported as .pin files which served as input for Mokapot.

### 2.4 Mokapot rescoring

The linear support vector machine (LSVM), which is also utilized by Percolator, serves as the default model in Mokapot (v0.10.0). This model, configured with default hyperparameters, is used as a baseline to which random forest^29^ and XGBoost^11^ models of varying complexity are compared. The features used to represent the PSMs are computed from the results of each one of the four search engines (Andromeda, Comet, MS-GF+, and MS-Amanda) and consist of the default MS^2^PIP^30^ and DeepLC^31^ features as outlined in the MS^2^Rescore^6^ publication’s Supplementary Table S2.

To create models with different complexities, we utilized the random forest^29^ learning algorithm from scikit-learn (v1.5.2)^32^. We varied the *max_depth* hyperparameter from 1 to 40 (values: 1, 2, 3, 4, 5, 6, 7, 8, 9, 10, 15, 20, 40) and the *min_samples_leaf* hyperparameter from 4 to 64 (values: 4, 8, 16, 32, 64) to generate models with different levels of complexity. The *max_depth* hyperparameter limits the number of sequential node-splitting decisions from root to leaf in a decision tree, while the *min_samples_leaf* hyperparameter specifies the minimum number of samples required at each leaf node. Increasing the *max_depth* value and decreasing the *min_samples_leaf* value results in a more complex random forest model. Systematically varying *max_depth* and *min_samples_leaf* across this range of values allows us to examine directly the relationship between model complexity and rescoring performance.

For completeness, we also evaluated the gradient boosting algorithm XGBoost (v1.7.6). Here, we followed the configuration outlined in the original Mokapot paper^5^, where hyperparameter optimization was performed via a Scikit-learn *GridSearchCV*, rather than by the systematic hyperparameters variation used for the random forest model.

To compare rescoring algorithms across datasets of different sizes, PSM counts were normalized by dividing the number of identified PSMs for each algorithm by the total number of PSMs for that dataset and multiplying by 100. The difference between these normalized values and the corresponding normalized Mokapot LSVM counts was then computed to determine the relative improvement or reduction in PSM identification.

### 2.5 Entrapment peptide sequences experiment

Utilizing more complex models for PSM rescoring carries the risk of overfitting to target PSMs. In such cases of overfitting, the likelihood of random matches aligning with the target peptide database increases compared to the decoy peptide database. Consequently, this can result in overly optimistic FDR estimates, as these are based on the assumption of equal matching probability between the target and decoy database for false matches.

To investigate this issue, we used an established entrapment procedure^33–35^. In this approach, the target peptide database is expanded by incorporation of entrapment peptides derived from nine randomly shuffled versions of the target proteome. The purpose of these entrapment peptides is to attract the majority of random (incorrect) target PSMs, thereby ensuring that the majority of PSMs aligning with the true target database are indeed correct. By enlarging the entrapment database, the probability of an assumed correct match in fact being true is increased. The entrapment FDR at a decoy FDR of 1% is computed as: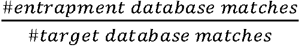

## 3. RESULTS

### 3.1 Assessing the effect of adding entrapment sequences to the search space

We first rescored the MS-GF+ search results obtained without extending the target database with entrapment sequences. This analysis was conducted exclusively for the MS-GF+ search engine and focused on three datasets (CD8 T cells, *Gammarus fossarum*, and HEK293 cells), as it was not the primary objective of this study but rather performed to provide a comparative baseline. These datasets were chosen to provide a diverse representation of experimental conditions and to enable a broader perspective on the performance across varying dataset sizes.

Rescoring results confirm that increasing the degree of complexity of the model used for semi-supervised rescoring increases the number of identified PSMs at 1% FDR (Supplementary Table 2). When comparing the results for the best-performing random forest model to the LSVM results, the number of identifications increases by 5.7% (CD8 T cells; from 64,059 to 67,740), 0.3% (*Gammarus fossarum*; from 206,887 to 207,546), and 0.2% (HEK293 cells; from 516,263 to 517,463). For the tuned XGBoost model, the number of identifications increases by 6% (CD8 T cells; from 64,059 to 67,889), 0.3% (*Gammarus fossarum*; from 206,887 to 207,653), and 0.45% (HEK293 cells; from 516,263 to 518,606).

We then rescored the MS-GF+ results obtained from a target database that contains both the target proteome and the entrapment sequences. As expected, this led to a reduction in identification sensitivity, due to the significantly expanded size of the search space (Supplementary Table 2). Nonetheless, the observed trend of increasing sensitivity with increasing model complexity remains consistent. Importantly, the entrapment FDR rises substantially with increasing model complexity, exceeding the 1% threshold for the more complex model, especially for the smaller dataset (Figure 1; Supplementary Table 2).

**Figure 1:**
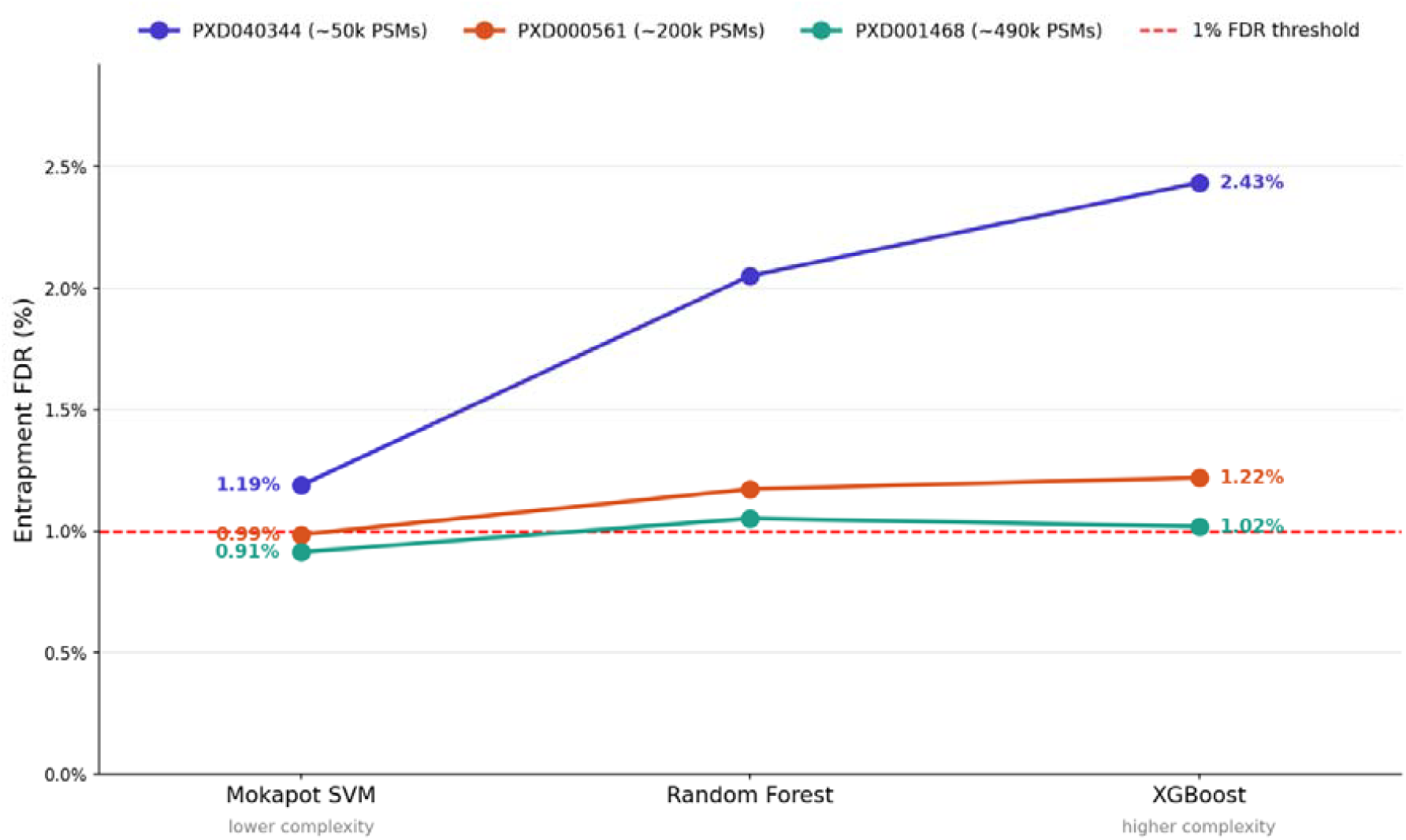
Entrapment FDR as a function of rescoring model complexity for MS-GF+ results across three datasets (PXD040344, PXD000561, and PXD001468). The x-axis orders the rescoring models by increasing complexity (Mokapot SVM, random forest, and XGBoost), while the y-axis shows the entrapment FDR (%). Each line corresponds to a dataset, with its approximate total number of identified PSMs noted in the legend, and each point is annotated with its entrapment FDR value. The dashed red line indicates the 1% entrapment FDR threshold.

### 3.2 Entrapment analysis across four search engines

In the next step, we extended the analysis to four search engines (Andromeda, Comet, MS-GF+, and MS Amanda), and rescored the results across eleven datasets, as listed in Supplementary Table 1. This analysis aimed to evaluate the impact of incorporating entrapment sequences on the number of identified PSMs. We start with baseline search engine results and progress through rescoring models of varying complexity. Only the results on PXD009815 are shown, for completeness the results of all other datasets are available in Supplementary figures 1 to 26.

#### 3.2.1 Baseline results for each search engine

The baseline results from the four search engines reveal the variability in identification sensitivity across datasets. Figure 2A illustrates these differences for dataset PXD009815. While not the main focus of this research, result show that MS-GF+ and Comet achieved higher numbers of identified PSMs on most datasets, while MS Amanda and Andromeda typically ranked third and fourth, respectively. For completeness, the results for all datasets are provided in Supplementary Figure 1.

**Figure 2:**
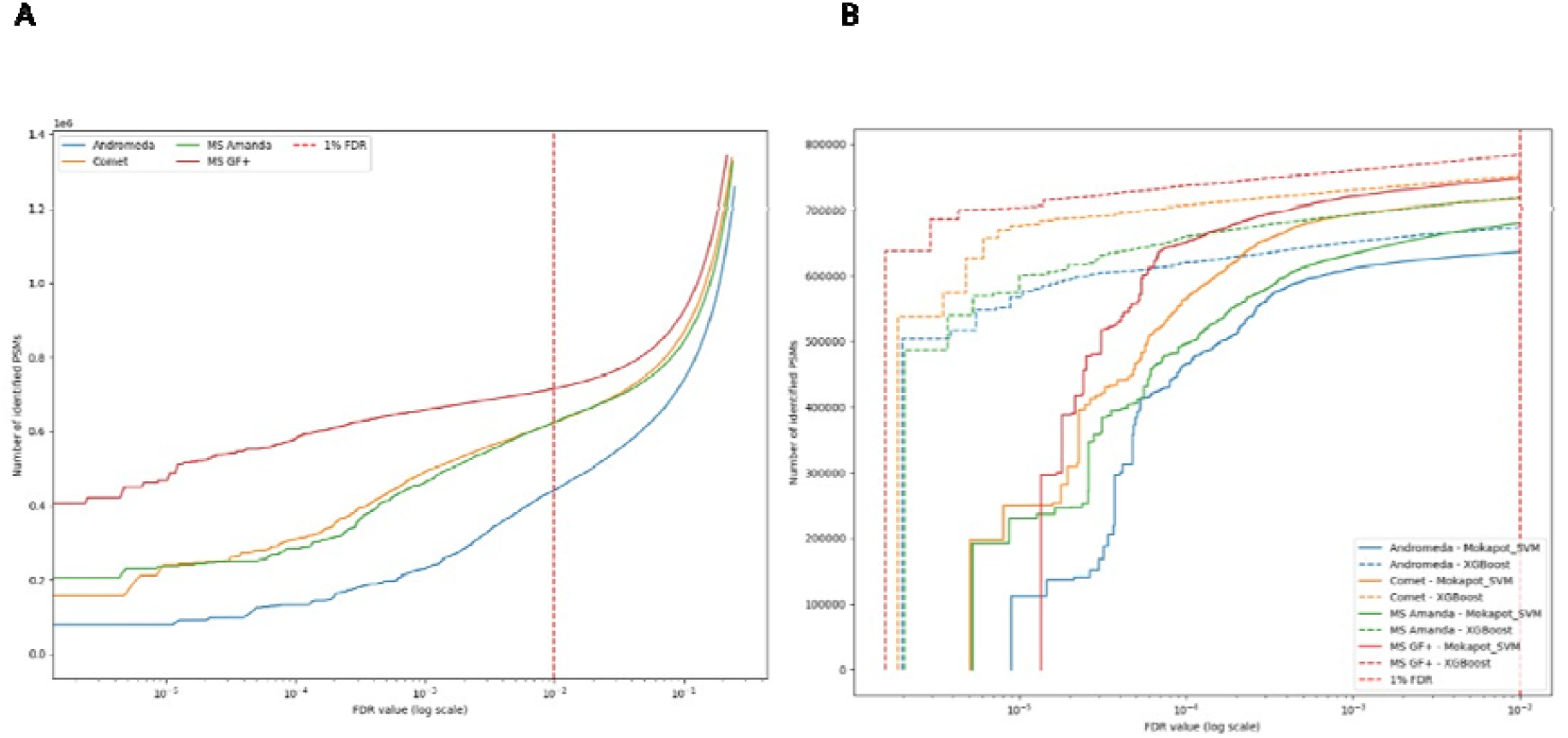
Comparison of peptide-spectrum match (PSM) identifications before and after rescoring for dataset PXD009815. (A) Cumulative numbers of PSMs identified without rescoring by Andromeda (blue), Comet (orange), MS Amanda (green), and MS-GF+ (red). (B) Cumulative numbers of PSMs identified after rescoring with mokapot using either a support vector machine (SVM; solid lines) or XGBoost (dashed lines). Colours indicate the corresponding search engines as in panel A. In both panels, the x-axis shows the false discovery rate (FDR) on a logarithmic scale, and the y-axis shows the cumulative number of identified PSMs. The vertical red dashed line marks the commonly used 1% FDR threshold.

#### 3.2.2 Rescoring with linear and non-linear algorithms in Mokapot for each search engine

Figure 2B presents the number of identified PSMs after rescoring with LSVM, default linear algorithm of Mokapot, and XGBoost, a non-linear algorithm, on a single dataset. The results show that the use of a non-linear algorithm can lead to an increase in the number of identified PSMs, regardless of the search engines used. However, this analysis reflects the total number of identified PSMs without providing insights into the ratio of target to entrapment identifications, requiring further investigation of these results. The outcomes for the other datasets are available in Supplementary Figure 2.

#### 3.2.3 Impact of Rescoring Algorithms on PSM Identification

To further investigate the differences in PSM counts after rescoring with different algorithms, we compared the results obtained from XGBoost and the most complex random forest model defined by *min_samples_leaf = 4 and max_depth = 40 (hereafter referred to as RandomForest_MSL4_MD40)*, against Mokapot LSVM across all datasets and search engines. Figure 3 visualizes these differences by plotting the difference in normalized PSM counts between each rescoring algorithm and Mokapot LSVM.

**Figure 3:**
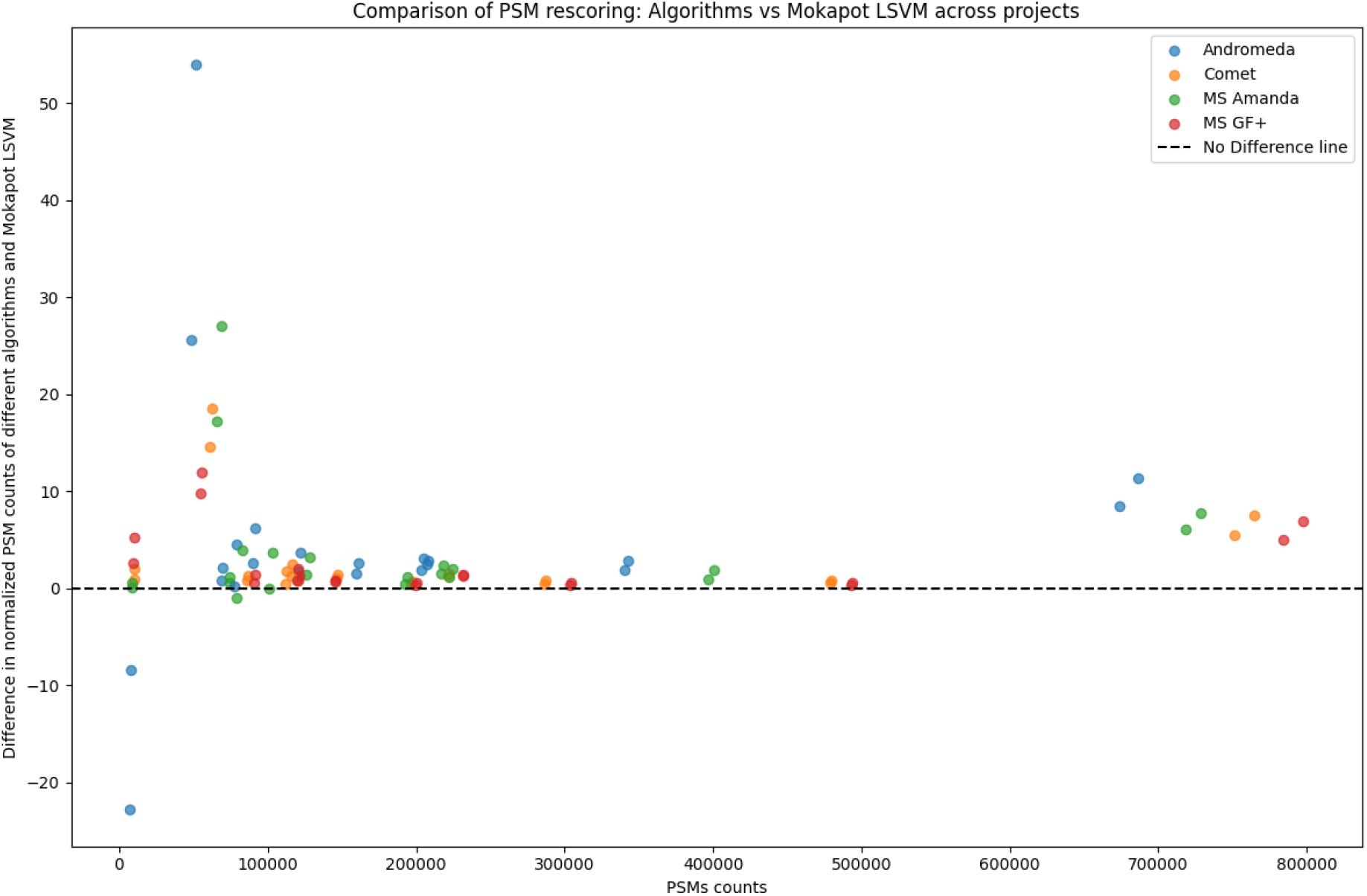
Comparison of PSM counts after rescoring with XGBoost and RandomForest_MSL4_MD40 relative to Mokapot LSVM across 11 datasets and four search engines. Each search engine-dataset combination is represented by two data points corresponding to XGBoost and RandomForest_MSL4_MD40. The x-axis represents the total number of identified PSMs, while the y-axis shows the difference in normalized PSM counts relative to Mokapot LSVM. Normalization was performed by dividing the PSM counts of each algorithm by the total number of PSMs in the dataset and multiplying by 100. The dashed horizontal line at y = 0 indicates no difference between Mokapot LSVM and the alternative rescoring algorithms. The results demonstrate that while XGBoost and RandomForest_MSL4_MD40 generally improve PSM counts, the effect is dataset and search engine dependent.

For each of the 11 datasets, results from four different search engines were considered. For each search engine, two data points are shown, corresponding to rescoring with XGBoost and *RandomForest_MSL4_MD40*. The x-axis represents the total number of identified PSMs, while the y-axis represents the difference in normalized PSM counts after rescoring. The dashed horizontal line at y = 0 serves as a reference, indicating no difference between the rescoring algorithms and Mokapot LSVM.

The results indicate that, in most cases, rescoring with XGBoost or *RandomForest_MSL4_MD40* leads to an increase in the number of identified PSMs compared to Mokapot LSVM rescoring. However, the extent of this improvement varies across datasets and search engines. The effect is particularly pronounced for certain search engines, while others only exhibit modest improvements or even reductions in sensitivity. Notably, two data points fall below the no difference line, indicating that more complex algorithms identified fewer PSMs than Mokapot LSVM. These cases correspond to datasets with small sample sizes.

#### 3.2.4 Considering the number of identified entrapment PSMs

To evaluate the accuracy of the observed increase in total PSMs, it is essential to also consider the changes in the number of identified entrapment PSMs. Figure 4 represents the number of identified target PSMs alongside the number of identified entrapment PSMs for different levels of model complexity for dataset PXD009815 searched with Comet. The corresponding figures for other search engines and datasets are available in Supplementary Figures 3 to 5.

**Figure 4:**
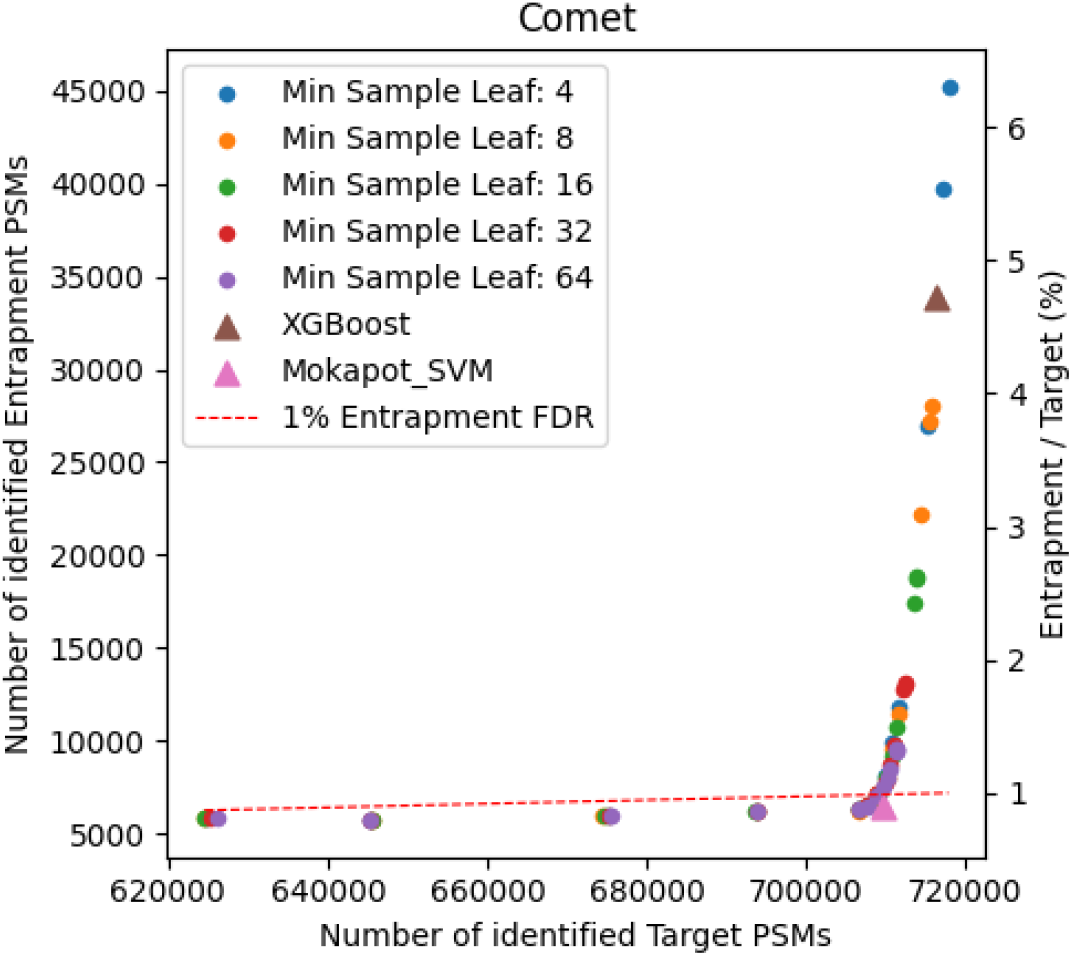
The number of identified target and entrapment PSMs for different model complexities when rescoring Comet search results for PXD009815. The x-axis shows the number of identified target PSMs and the left y-axis shows the number of identified entrapment PSMs, while the right y-axis reports the corresponding entrapment-to-target ratio as a percentage. Each point represents a rescoring model, with varying minimum sample leaf values for the random forest models (colored circles), alongside results for XGBoost (brown triangle) and Mokapot SVM (pink triangle). For each minimum sample leaf value, 13 different maximum depth settings were also evaluated. The dashed red line indicates the 1% entrapment FDR threshold.

Figure 4 demonstrates that as the complexity of the random forest model increases, the results closely follow the 1% entrapment FDR line, up to a point where the number of entrapment matches starts to increase exponentially. This point is near the LSVM model complexity. An extreme example is the XGBoost model and the most complex random forest models, which identify roughly three times as many entrapment PSMs as the LSVM model, despite achieving only a modest increase in the number of target PSMs. The entrapment FDR for the optimized XGBoost model is 5%, and for most complex random forest model is 6.3%.

#### 3.2.5 Heatmap analysis of random forest performance

To further investigate the impact of random forest rescoring, we generated heatmaps to visualize the distribution of identified target PSMs, annotated by the entrapment FDR. With the decoy FDR ratio set to 1%, we expect the entrapment FDR ratio to also approximate this value. However, this is not always the case, particularly for more complex models, where the entrapment FDR ratio often exceeds 1%.

We calculated the entrapment FDR ratio for various hyperparameters of the random forest model, as shown in Figure 5 for dataset PXD009815 searched with Comet. In this figure, areas with a higher number of identified PSMs are represented in er colors, simultaneously annotated with higher entrapment FDR values. The area highlighted by blue dotted rectangle indicates the optimal combination of hyperparameters for this specific dataset, as computed by our script which chooses the highest number of target PSMs, while maintaining an entrapment FDR closest to the desired value of 1%. Additional figures for other datasets and search engines are provided in Supplementary Figures 6 to 8.

**Figure 5:**
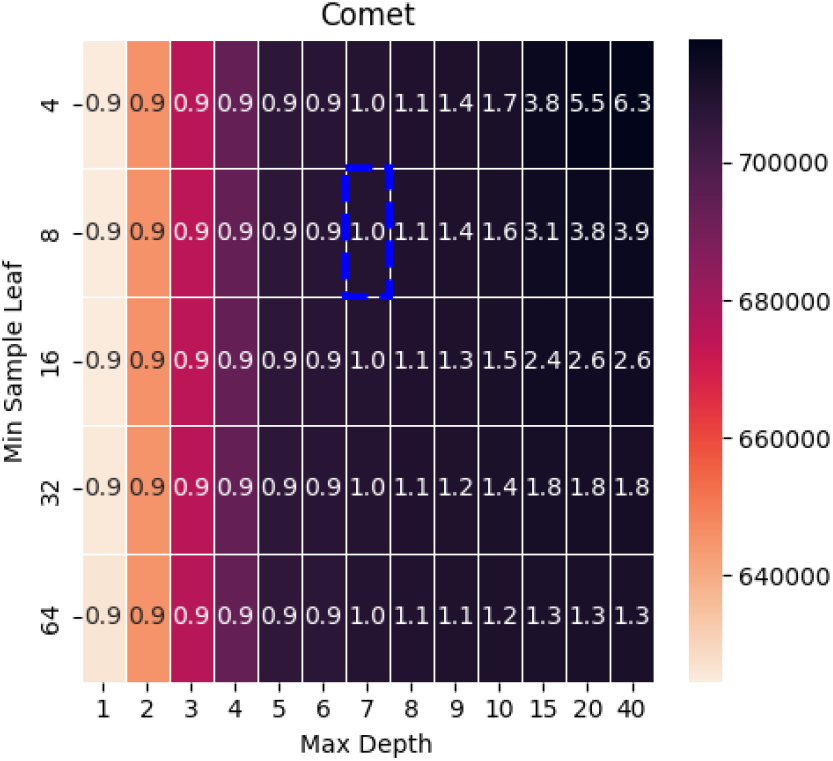
Heatmap showing the distribution of identified target PSMs across different hyperparameter settings for the random forest model when rescoring Comet search results for dataset PXD009815. The x-axis represents the maximum tree depth, while the y-axis corresponds to the minimum sample leaf parameter. Darker colors indicate a higher number of identified target PSMs, with each cell annotated by the corresponding entrapment FDR value. The dashed blue rectangle highlights the optimal hyperparameter region, where the highest number of target PSMs is achieved while maintaining an entrapment FDR closest to 1%.

#### 3.2.6 UpSet plot analysis for target and entrapment PSMs

Finally, UpSet plots were used to visualize the overlap and differences in identified PSMs among different search engines and rescoring algorithms. Figure 6 illustrates the intersection of target (Figure 6A) and entrapment (Figure 6B) PSM identifications across different search engines rescored by XGBoost. The plot reveals that entrapment PSMs were largely unique to each search engine, indicating that most identifications were random. This uniqueness reflects the random nature of the entrapment sequences, which lack any biological identification bias and therefore produce no consistent, reproducible matches across search engines. In contrast, target PSMs demonstrated a high degree of agreement across search engines.

**Figure 6:**
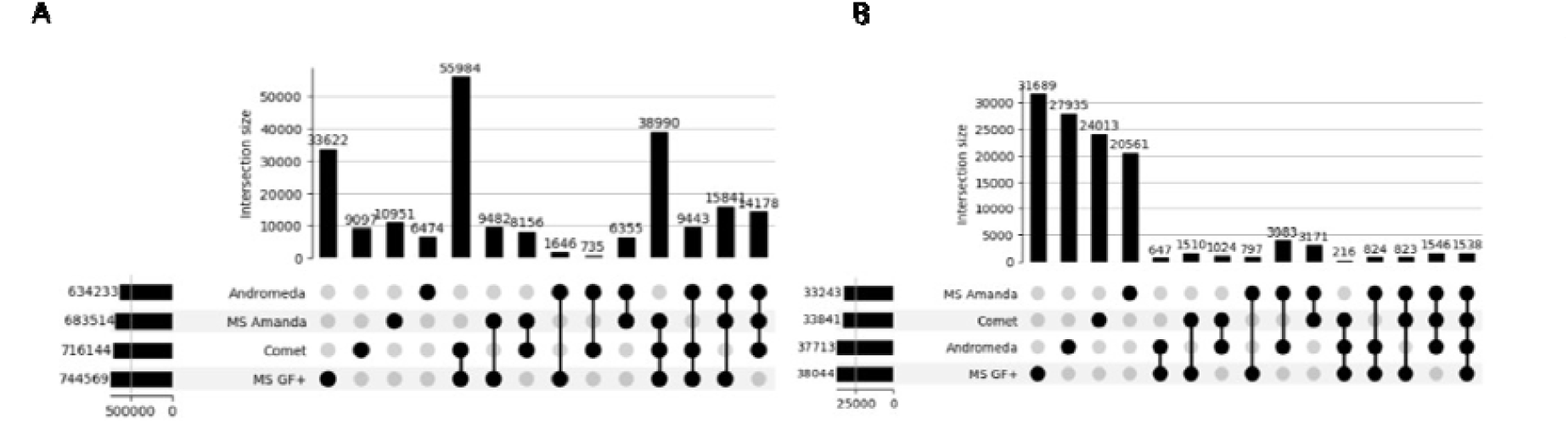
Overlap of peptide-spectrum match (PSM) identifications across search engines after XGBoost-based rescoring for dataset PXD009815. UpSet plots show intersections among PSMs identified by the different search engines for (A) target PSMs and (B) entrapment PSMs. The horizontal axis represents unique and shared intersections among search engines, whereas the vertical axis indicates the number of PSMs in each intersection. Target PSM identifications showed substantial overlap across search engines, while entrapment PSMs were predominantly unique to individual search engines, consistent with a largely random distribution of entrapment matches. For panel A, the intersection shared by all search engines, comprising 579,561 PSMs, was omitted to improve visualization of the remaining intersections.

For the target PSMs figure, we excluded the intersection of all search engines, with a value of 579,561, as it represented the highest peak, making the other peaks easier to compare. These results align with our expectation that random identifications (i.e., “lucky guesses”) for target PSMs would be minimized. Additional figures for datasets are provided in Supplementary Figures 9 to 14.

In Figure 7, the intersection of target and entrapment PSM identifications for different algorithms searched with Comet is presented. Similar to Figure 6, entrapment PSMs were largely unique to each algorithm, but there was greater overlap among certain algorithms, such as XGBoost and more complex random forest models. This can be explained by the fact that we rescored results from a single search engine, which increases the likelihood of shared entrapment PSMs compared to the results from different search engines.

**Figure 7:**
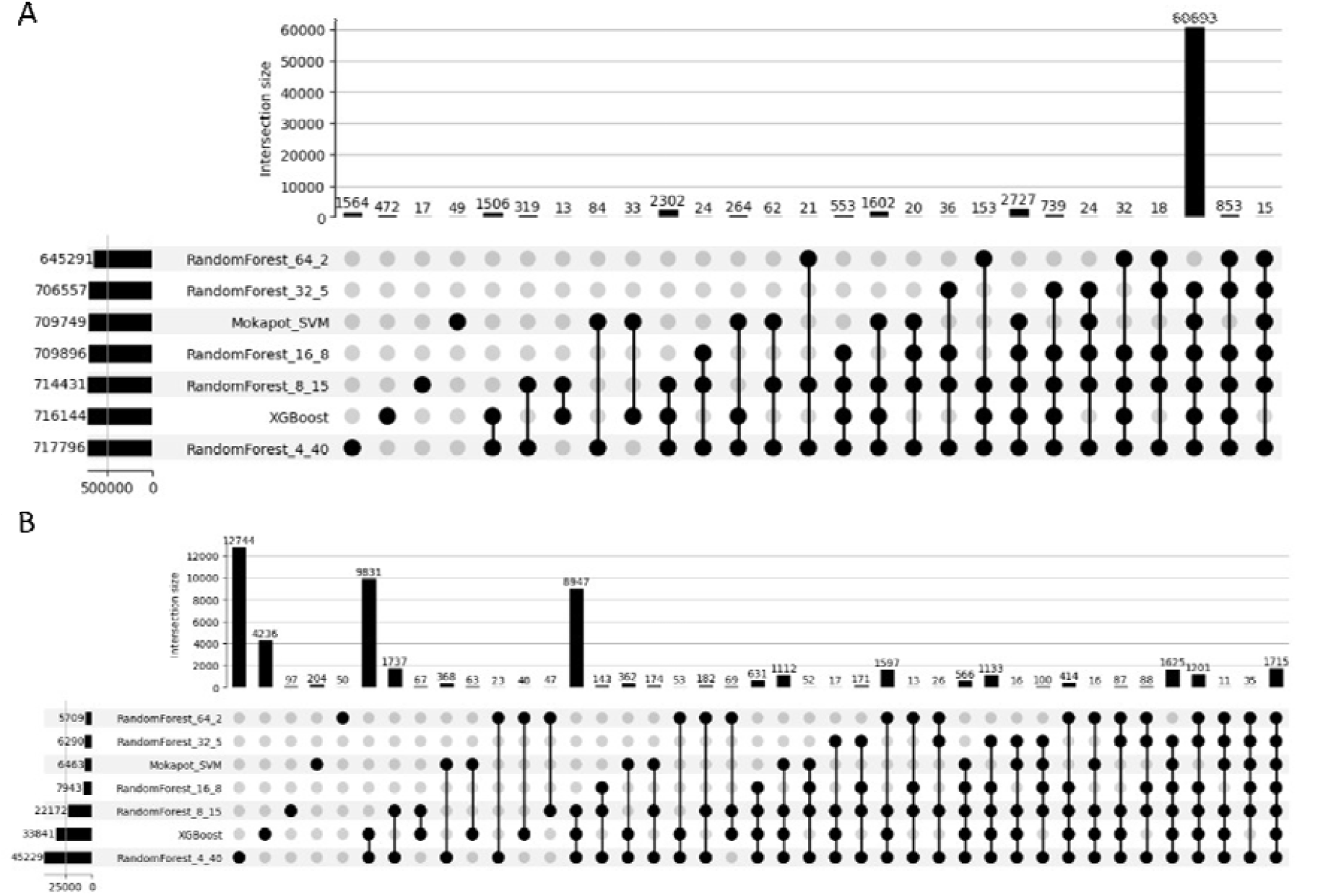
Overlap of peptide-spectrum match (PSM) identifications across rescoring algorithms applied to Comet search results for dataset PXD009815. UpSet plots show the intersections of PSMs identified by the different rescoring methods for (A) target PSMs and (B) entrapment PSMs. The horizontal axis represents unique and shared intersections among the rescoring algorithms, whereas the vertical axis indicates the number of PSMs in each intersection. In panel A, the intersection shared by all rescoring algorithms, comprising 644,148 target PSMs, was omitted to improve visualization of the remaining intersections.

For the target PSMs figure, the intersection of all algorithms has a value of 644,148. As expected, the second highest peak corresponds to the intersection of all algorithms excluding the least complex one. Additional figures for different search engines and datasets are provided in Supplementary Figures 15 to 26.

## 4 Discussion and Conclusion

We examined the use of varying degrees of model complexity to rescore PSMs obtained from four different search engines (Andromeda, Comet, MS-GF+, and MS Amanda). We conducted searches on eleven diverse MS/MS spectrum files against different target databases, comprising the target proteome with and without the addition of entrapment peptide sequences.

The results show that rescoring models with increasing complexity can effectively increase the number of identified PSMs across all datasets. However, this increase in identifications comes at the cost of many false positive identifications, which are not properly controlled with the standard target-decoy approach. This increase in false identifications means that the use of excessively complex models in PSM rescoring introduces a bias towards favoring random matches from the target proteome database, thereby leading to overly optimistic FDR estimations.

From the results it is also clear that an entrapment strategy can be used to estimate the level of overfitting in the rescoring model. This measure of overfitting can be used to select the level of complexity necessary to achieve both high sensitivity and properly controlled PSM FDR. However, the computational cost of searching with an entrapment database is high and likely impractical for most analyses. Due to this limitation, further research is needed to determine the level of complexity of a rescoring model that ensures sensitivity without overfitting.

## 5 Data availability

All data and script used to make the comparison and to generate figures are available on Zenodo at https://doi.org/10.5281/zenodo.15535864.

## 6 Author contributions

Alireza Nameni: Conceptualization, Methodology, Formal analysis, Writing – original draft, Writing – review & editing Arthur Declercq: Software, Writing – review & editing Ralf Gabriels: Software, Writing – review & editing Sven Degroeve: Conceptualization, Methodology, Writing – review & editing Lennart Martens: Writing – review & editing Robbin Bouwmeester: Conceptualization, Methodology, Formal analysis, Supervision, Validation, Writing – original draft, Writing – review & editing

## 7 Acknowledgments

A.N. acknowledges funding from the European Union’s Horizon 2020 research and innovation programme under the Marie Skłodowska-Curie grant agreement N° 956148. S.D. acknowledges funding from the European Union’s Horizon 2020 Programme grant no. H2020-INFRAIA-2018-1. A.D., R.G., L.M. and R.B. acknowledge funding from the Research Foundation Flanders (FWO) [1SE3724N 12AK526N, G010023N, G028821N, 12A6L24N]. L.M. acknowledges funding from the Horizon Europe Projects BAXERNA 2.0 [101080544] and COMBINE [101191739], and from the Ghent University Concerted Research Action [BOF21/GOA/033]. L.M. is further supported by the CHIST-ERA project ODEEP-EU [G0GDV23N].

