## Supplementary document for "Evaluating the use of non-linear models in data-driven rescoring of peptide-spectrum matches"

**Table of Contents**

| Table S1 | List of PRIDE projects used to conduct evaluation, with raw files, instrument, sample, and FASTA file |
| --- | --- |
| Table S2 | Mokapot rescoring results |
| Figure S1 | Comparison of identified PSMs before any rescoring across four search engines |
| Figure S2 | Comparison of identified PSMs after rescoring with Mokapot SVM and XGBoost across four search engines |
| Figure S3–S5 | Target vs. entrapment PSMs for different model complexities, across all datasets |
| Figure S6–S8 | Heatmaps of identified target PSMs across hyperparameter settings, across all datasets |
| Figure S9–S14 | UpSet plots of identified PSMs across search engines after XGBoost rescoring, across all datasets |
| Figure S15–S20 | UpSet plots of identified PSMs across rescoring algorithms for Comet, across all datasets |
| Figure S21–S26 | UpSet plots of identified PSMs across rescoring algorithms for MS-GF+, across all datasets |

Supplementary Table 1: List of PRIDE projects used to conduct evaluation of our experiment and their corresponding used Raw files, along with the instrument, sample and the corresponding FASTA file.

| PRIDE project | Raw data files used | Sample and Instrument | FASTA file |
| --- | --- | --- | --- |
| PXD000561 | Adult_CD8Tcells_Gel_Elite_44_f01  Adult_CD8Tcells_Gel_Elite_44_f02  Adult_CD8Tcells_Gel_Elite_44_f03  Adult_CD8Tcells_Gel_Elite_44_f04  Adult_CD8Tcells_Gel_Elite_44_f05  Adult_CD8Tcells_Gel_Elite_44_f06  Adult_CD8Tcells_Gel_Elite_44_f07  Adult_CD8Tcells_Gel_Elite_44_f08  Adult_CD8Tcells_Gel_Elite_44_f09  Adult_CD8Tcells_Gel_Elite_44_f10  Adult_CD8Tcells_Gel_Elite_44_f11  Adult_CD8Tcells_Gel_Elite_44_f12  Adult_CD8Tcells_Gel_Elite_44_f13  Adult_CD8Tcells_Gel_Elite_44_f14  Adult_CD8Tcells_Gel_Elite_44_f15  Adult_CD8Tcells_Gel_Elite_44_f16  Adult_CD8Tcells_Gel_Elite_44_f17  Adult_CD8Tcells_Gel_Elite_44_f18  Adult_CD8Tcells_Gel_Elite_44_f19  Adult_CD8Tcells_Gel_Elite_44_f20  Adult_CD8Tcells_Gel_Elite_44_f21  Adult_CD8Tcells_Gel_Elite_44_f22  Adult_CD8Tcells_Gel_Elite_44_f23  Adult_CD8Tcells_Gel_Elite_44_f24 | CD8 T cells (LTQ Orbitrap Velos Elite) | uniprotHomoReview20365+contaminants |
| PXD006675 | 20160901_QEp2_SoDo_SA_LC12-13_PV8-frac1  20160901_QEp2_SoDo_SA_LC12-13_PV8-frac2  20160901_QEp2_SoDo_SA_LC12-13_PV8-frac3  20160901_QEp2_SoDo_SA_LC12-13_PV8-frac4  20160901_QEp2_SoDo_SA_LC12-13_PV8-frac5  20160901_QEp2_SoDo_SA_LC12-13_PV8-frac6  20160901_QEp2_SoDo_SA_LC12-13_PV8-frac7  20160901_QEp2_SoDo_SA_LC12-13_PV8-frac8_160906090046 | Human heart tissue (Q Exactive) | uniprotHomoReview20365+contaminants |
| PXD001468 | b1906_293T_proteinID_01A_QE3_122212  b1922_293T_proteinID_02A_QE3_122212  b1922_293T_proteinID_03A_QE3_122212  b1922_293T_proteinID_04A_QE3_122212  b1925_293T_proteinID_05A_QE3_122212  b1926_293T_proteinID_06A_QE3_122212  b1927_293T_proteinID_07A_QE3_122212  b1928_293T_proteinID_08A_QE3_122212  b1929_293T_proteinID_09A_QE3_122212  b1930_293T_proteinID_10A_QE3_122212  b1931_293T_proteinID_11A_QE3_122212  b1932_293T_proteinID_12A_QE3_122212  b1906_293T_proteinID_01B_QE3_122212  b1922_293T_proteinID_02B_QE3_122212  b1922_293T_proteinID_03B_QE3_122212  b1922_293T_proteinID_04B_QE3_122212  b1925_293T_proteinID_05B_QE3_122212  b1926_293T_proteinID_06B_QE3_122212  b1927_293T_proteinID_07B_QE3_122212  b1928_293T_proteinID_08B_QE3_122212  b1929_293T_proteinID_09B_QE3_122212  b1930_293T_proteinID_10B_QE3_122212  b1931_293T_proteinID_11B_QE3_122212  b1932_293T_proteinID_12B_QE3_122212 | HEK293 cells (Q Exactive) | uniprotHomoReview20365+contaminants |
| PXD001250 | 20140711_EXQ00_KiSh_SA_Brain_1  20140711_EXQ00_KiSh_SA_Brain_2  20140711_EXQ00_KiSh_SA_Brain_3  20140711_EXQ00_KiSh_SA_Brain_4 | Mouse brain tissue (Q Exactive) | UP000000589_10090_Mouse |
| PXD000612 | 20120329_EXQ5_KiSh_SA_LabelFree_HeLa_Proteome_PV_rep1_pH3  20120329_EXQ5_KiSh_SA_LabelFree_HeLa_Proteome_PV_rep1_pH4  20120329_EXQ5_KiSh_SA_LabelFree_HeLa_Proteome_PV_rep1_pH5  20120329_EXQ5_KiSh_SA_LabelFree_HeLa_Proteome_PV_rep1_pH6  20120329_EXQ5_KiSh_SA_LabelFree_HeLa_Proteome_PV_rep1_pH8  20120329_EXQ5_KiSh_SA_LabelFree_HeLa_Proteome_PV_rep1_pH11 | HeLa cells (Q Exactive) | uniprotHomoReview20365+contaminants |
| PXD004948 | 20160323_CoAN_CTRL1_3372  20160323_CoAN_CTRL2_3373  20160329_CoAN_CTRL3_DR_3406 | Mouse (Q Exactive) | UP000000589_10090_Mouse |
| PXD040344 | E00377_MS21-022_Male_1_Caecum-4  E00377_MS21-022_Male_1_Caecum-5  E00377_MS21-022_Male_1_Caecum-6 | Gills and caeca from *Gammarus fossarum* (Q Exactive) | T-GFBM_filtered_orf_GHDA01.1_2021-06-21 |
| PXD004947 | 03022016_Clara_MP_Fraction_02  03022016_Clara_MP_Fraction_03  03022016_Clara_MP_Fraction_04  03022016_Clara_MP_Fraction_05  03022016_Clara_MP_Fraction_06  03022016_Clara_MP_Fraction_07  03022016_Clara_MP_Fraction_08  03022016_Clara_MP_Fraction_09  03022016_Clara_MP_Fraction_10  03022016_Clara_MP_Fraction_11  03022016_Clara_MP_Fraction_12  03022016_Clara_MP_Fraction_13  03022016_Clara_MP_Fraction_14  03022016_Clara_MP_Fraction_15  03022016_Clara_MP_Fraction_16  03022016_Clara_MP_Fraction_17  03022016_Clara_MP_Fraction_18  03022016_Clara_MP_Fraction_19  03022016_Clara_MP_Fraction_20  03022016_Clara_MP_Fraction_21  03022016_Clara_MP_Fraction_22  03022016_Clara_MP_Fraction_23  03022016_Clara_MP_Fraction_24  03022016_Clara_MP_Fraction_25  03022016_Clara_MP_Fraction_26  03022016_Clara_MP_Fraction_27  03022016_Clara_MP_Fraction_28  03022016_Clara_MP_Fraction_29  03022016_Clara_MP_Fraction_30  03022016_Clara_MP_Fraction_31 | *Solanum lycopersicum* (Q Exactive) | UP000004994_4081 |
| PXD004565 | 150710_QEp_PK_Bsub_DG_Br1  150710_QEp_PK_Bsub_DG_Br2  150710_QEp_PK_Bsub_DG_Br3  150710_QEp_PK_Bsub_DG_Br4 | *Bacillus subtilis* (Q Exactive) | UP000001570_224308 |
| PXD004325 | Mm2DLC_N_1_01  Mm2DLC_N_1_02  Mm2DLC_N_1_03  Mm2DLC_N_1_04  Mm2DLC_N_1_05  Mm2DLC_N_1_06  Mm2DLC_N_1_07  Mm2DLC_N_1_08  Mm2DLC_N_1_09  Mm2DLC_N_1_10  Mm2DLC_N_1_11  Mm2DLC_N_1_12 | *Methanosarcina mazei* (Q Exactive) | 91_methanosarcina_sp_crap_peptides_updated |
| PXD009815 | QEKAC160601_02  QEKAC160601_04  QEKAC160601_06  QEKAC160601_08  QEKAC160601_10  QEKAC160601_12  QEKAC160601_14  QEKAC160601_16  QEKAC160601_18  QEKAC160601_20  QEKAC160601_31  QEKAC160601_34  QEKAC160601_36  QEKAC160601_38  QEKAC160601_40  QEKAC160601_42  QEKAC160601_44  QEKAC160601_46  QEKAC160601_48  QEKAC160601_50  QEKAC160601_61  QEKAC160601_63  QEKAC160601_65  QEKAC160601_69  QEKAC160601_71  QEKAC160601_73  QEKAC160601_75  QEKAC160601_77  QEKAC160601_79  QEKAC160601_81  QEKAC160601_113  QEKAC160601_115  QEKAC160601_117  QEKAC160601_119  QEKAC160601_121  QEKAC160601_123  QEKAC160601_125  QEKAC160601_127  QEKAC160601_129  QEKAC160601_131 | *Saccharomyces cerevisiae* (Q Exactive) | uniprot_contaminant_yeast_ups_prot_03022023 |

Supplementary Table 2: Mokapot rescoring results. The model used for semi-supervised rescoring is listed in column algorithm where RF_MSL4_MD40refers to the Random Forest model with min sample leaf of 4 and max depth of 40. The number of identified PSMs in the search without entrapment peptides in the target database is listed in column total PSMs (no entrapment). The number of identified PSMs in the search with entrapment peptides in the target database is listed in column total PSMs (with entrapment). The entrapment FDR that corresponds to the total PSMs (with entrapment) column are listed in column entrapment-FDR (%).

| dataset | algorithm | Total PSMs (no entrapment) | Total PSMs (with entrapment) | entrapment-FDR (%) |
| --- | --- | --- | --- | --- |
| CD8 T cells | LSVM | 64059 | 51523 | 1.2 |
|  | RF_ MSL4_MD40 | 67740 | 54974 | 2.1 |
|  | XGBoost | 67889 | 55737 | 2.4 |
| *Gammarus fossarum* | LSVM | 206887 | 199064 | 1 |
|  | RF_ MSL4_MD40 | 207546 | 199794 | 1.2 |
|  | XGBoost | 207653 | 200059 | 1.2 |
| HEK239 cells | LSVM | 516263 | 491417 | 0.9 |
|  | RF_ MSL4_MD40 | 517463 | 492955 | 1.1 |
|  | XGBoost | 518606 | 493779 | 1 |

| PXD000561 | PXD006675 | PXD001468 |
| --- | --- | --- |
| 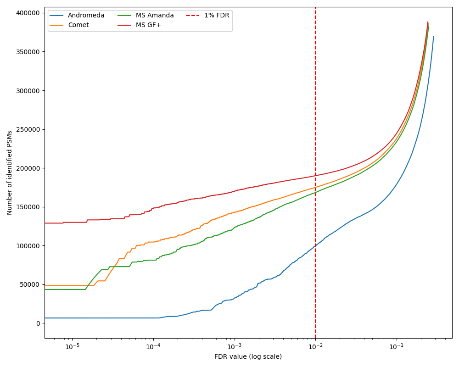 | 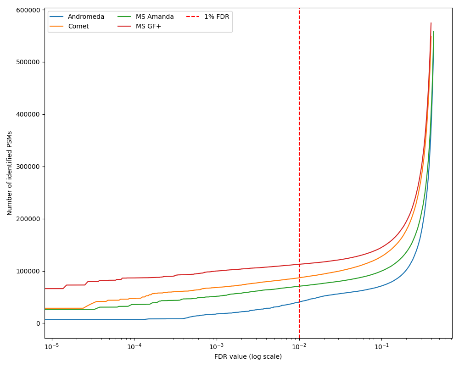 | 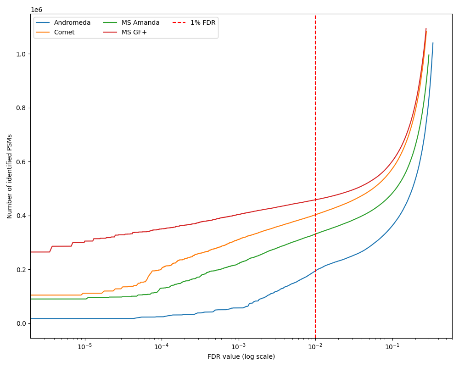 |
| PXD001250 | PXD000612 | PXD004948 |
| 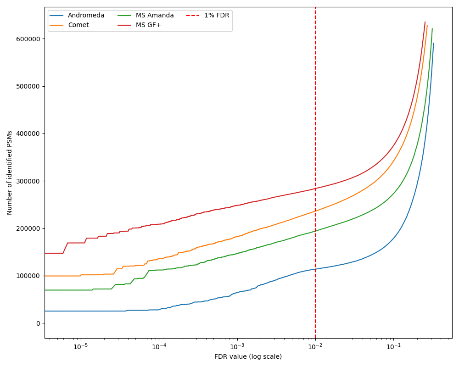 | 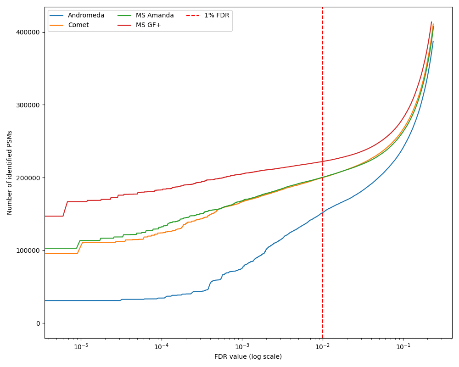 | 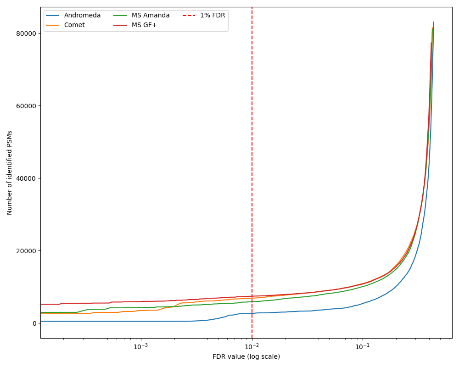 |
| PXD040344 | PXD004947 | PXD004565 |
| 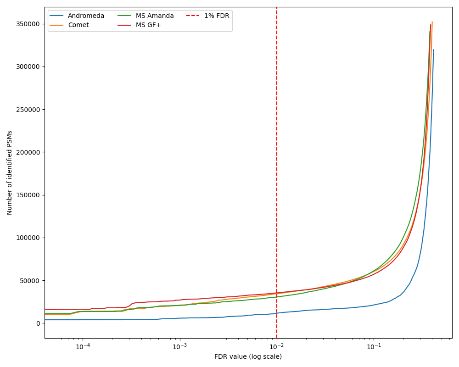 | 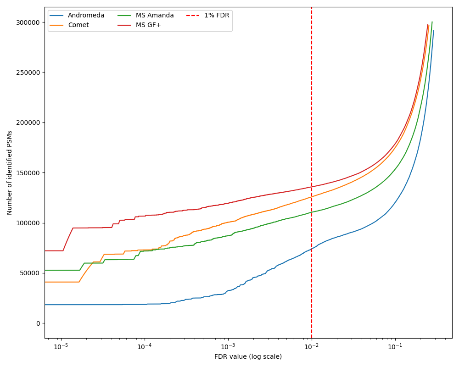 | 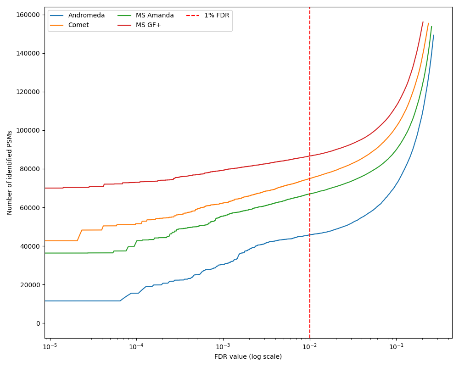 |
| PXD004325 | PXD009815 |  |
| 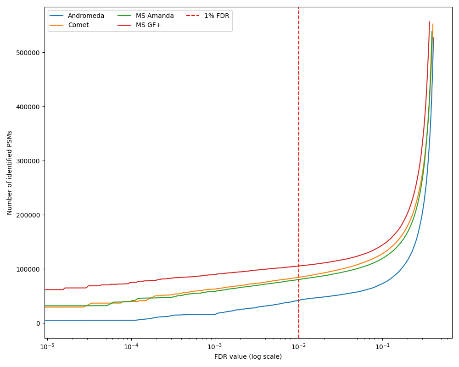 | 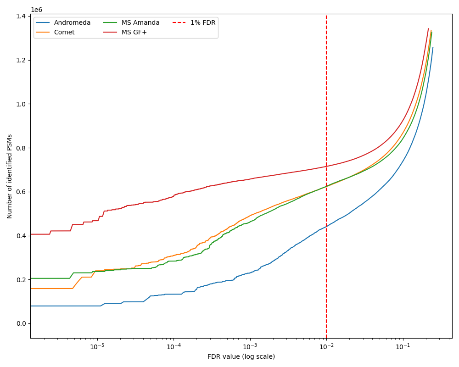 |  |

Supplementary Figure 1: Comparison of identified PSMs before any rescoring across four search engines for all analyzed datasets: Andromeda (blue), Comet (orange), MS Amanda (green), and MS-GF+ (red). The x-axis represents the FDR on a logarithmic scale, while the y-axis shows the cumulative number of identified PSMs. A vertical dashed red line indicates the commonly used 1% FDR threshold.

| PXD000561 | PXD006675 | PXD001468 |
| --- | --- | --- |
| 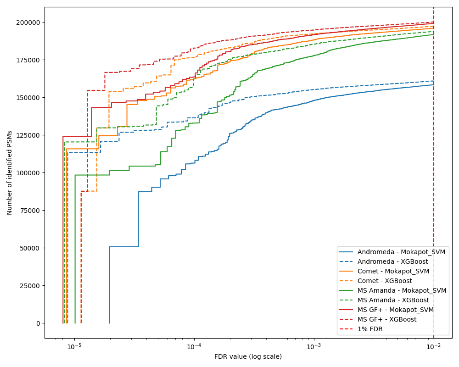 | 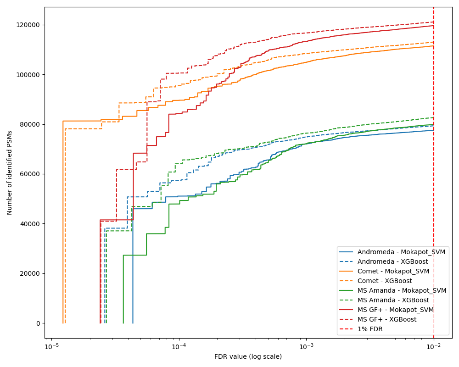 | 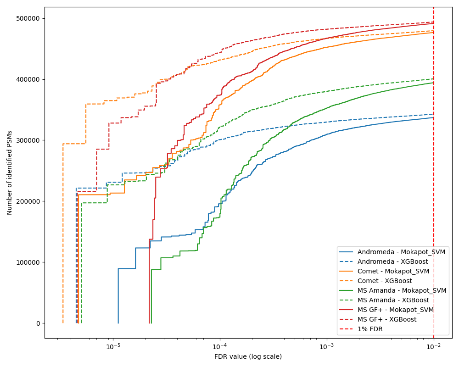 |
| PXD001250 | PXD000612 | PXD004948 |
| 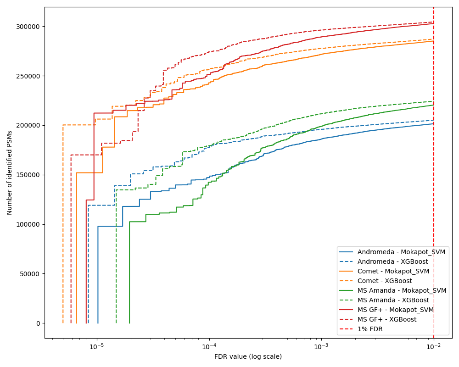 | 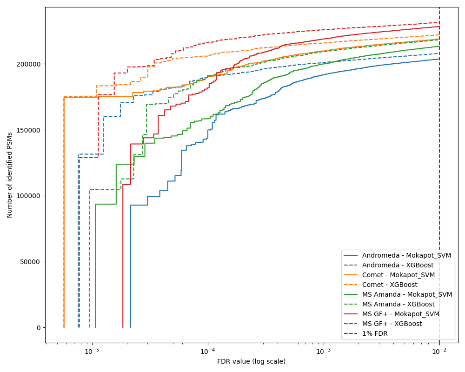 | 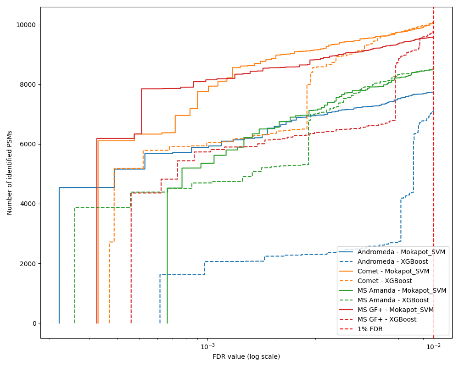 |
| PXD040344 | PXD004947 | PXD004565 |
| 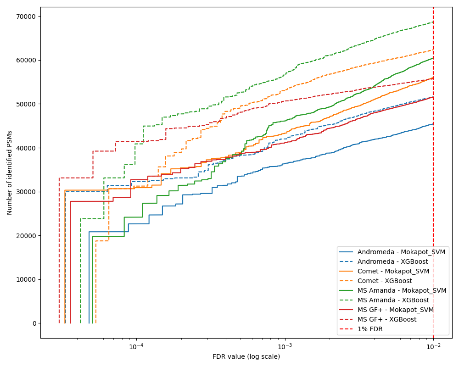 | 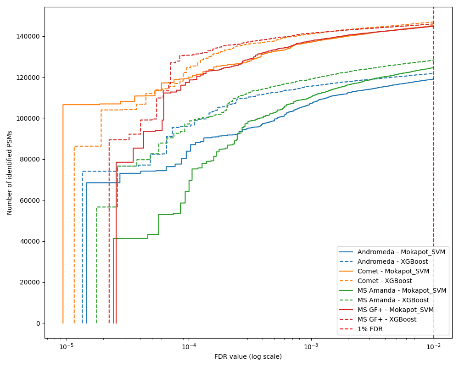 | 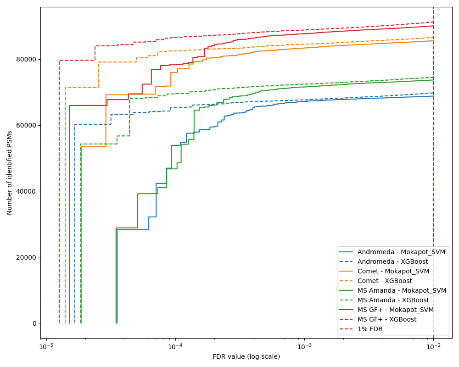 |
| PXD004325 | PXD009815 |  |
| 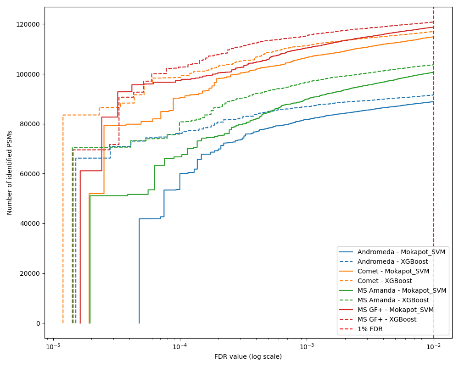 | 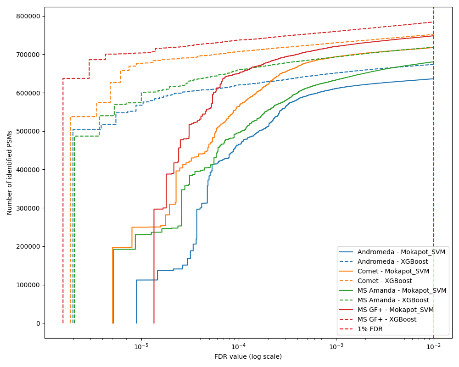 |  |

Supplementary Figure 2: Comparison of identified PSMs after rescoring with Mokapot SVM (solid lines) and XGBoost (dashed lines) across four search engines for all analyzed datasets: Andromeda (blue), Comet (orange), MS Amanda (green), and MS-GF+ (red). The x-axis represents the FDR on a logarithmic scale, while the y-axis shows the cumulative number of identified PSMs. A vertical dashed red line indicates the commonly used 1% FDR threshold.

PXD000561

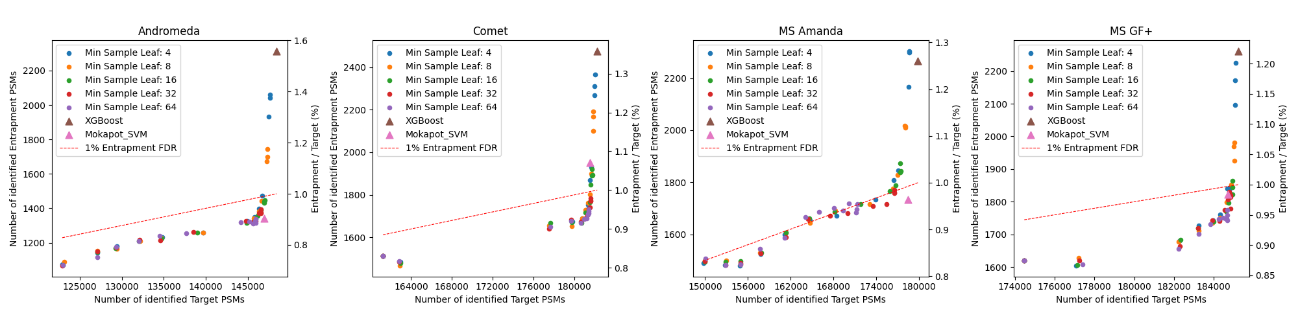

PXD006675

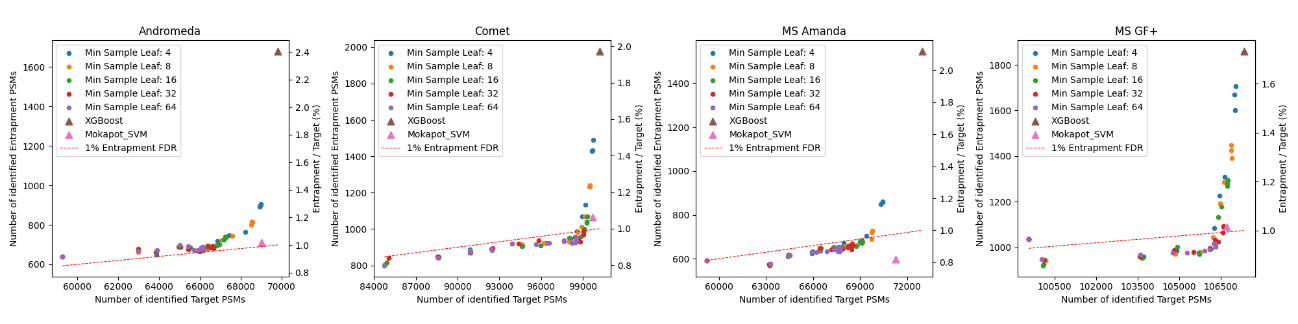

PXD001468

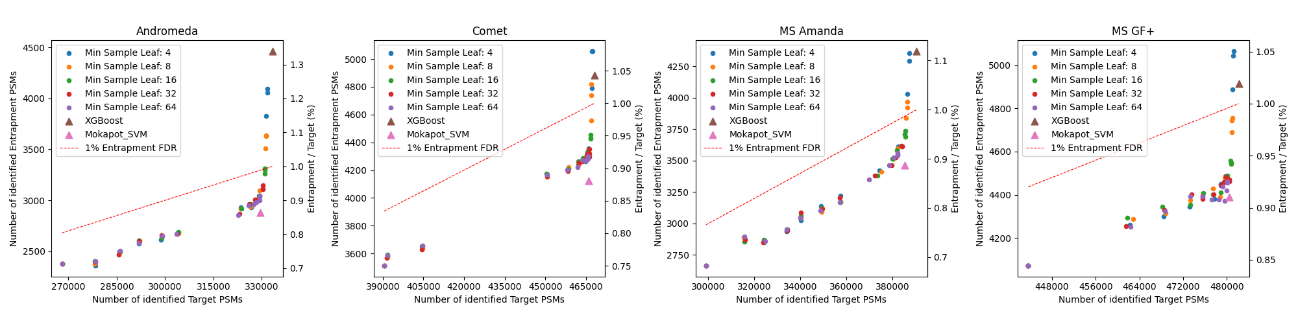

PXD001250

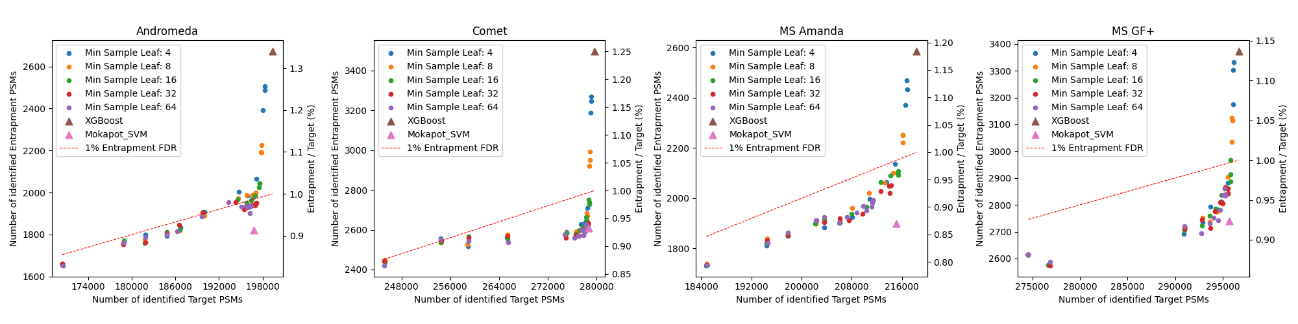

Supplementary Figure 3: Relationship between the number of identified target and entrapment PSMs for different model complexities when rescoring search results for datasets PXD000561, PXD006675, PXD001468, and PXD001250. Each panel corresponds to a different search engine (Andromeda, Comet, MS Amanda, and MS-GF+), displaying the distribution of identified PSMs for various rescoring models. Each point represents a model with different minimum sample leaf values for the random forest models (colored circles), alongside results for XGBoost (brown triangle) and Mokapot SVM (pink triangle). For each minimum sample leaf value, 13 different maximum depth settings were evaluated. The dashed red line indicates the 1% entrapment FDR threshold. Across all search engines and datasets, increasing model complexity generally results in a higher number of entrapment PSMs, particularly for highly complex random forest models and XGBoost.

PXD000612

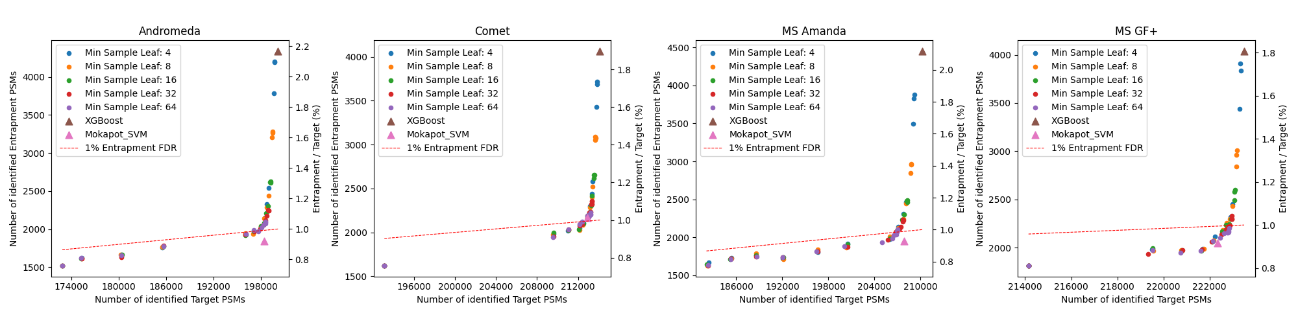

PXD004948

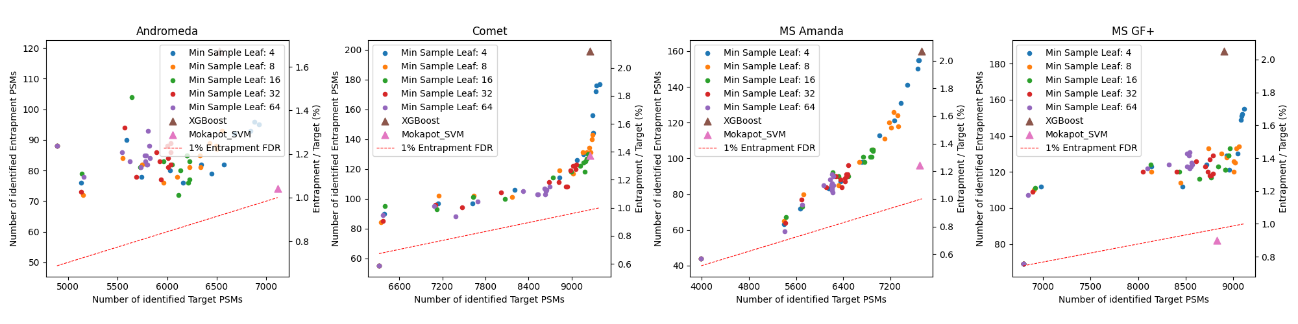

PXD040344

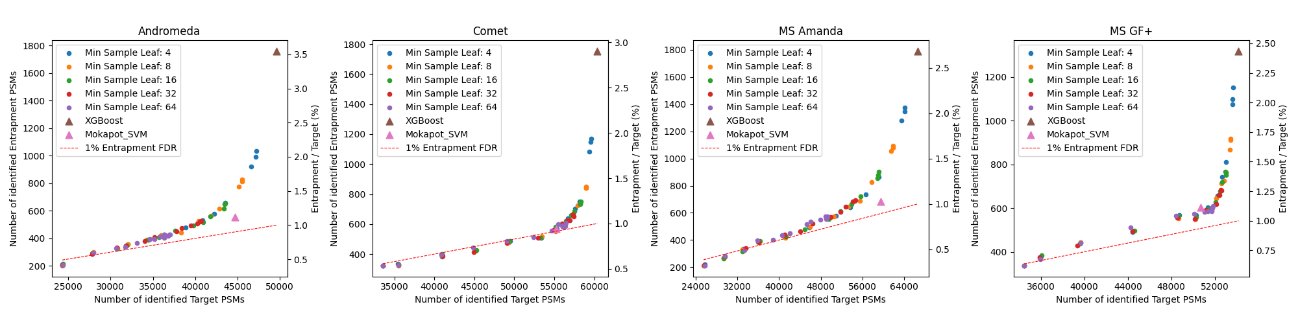

PXD004947

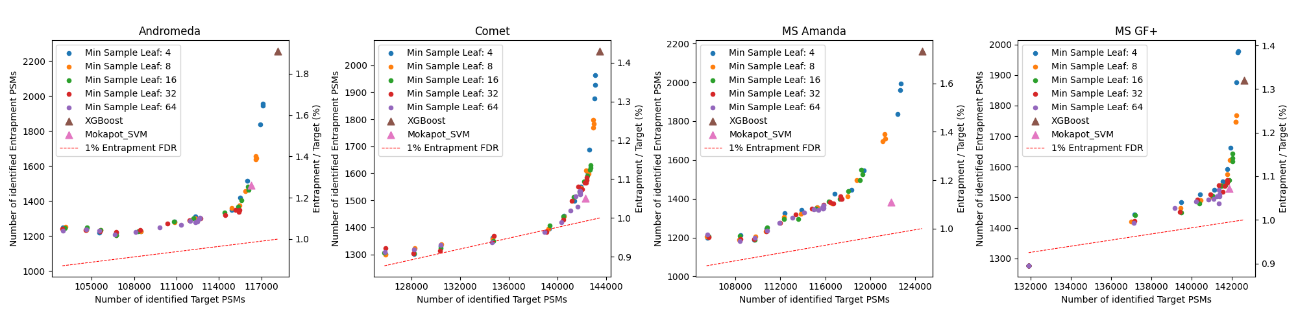

Supplementary Figure 4: Relationship between the number of identified target and entrapment PSMs for different model complexities when rescoring search results for datasets PXD000612, PXD004948, PXD040344, and PXD004947. Each panel corresponds to a different search engine (Andromeda, Comet, MS Amanda, and MS-GF+), displaying the distribution of identified PSMs for various rescoring models. Each point represents a model with different minimum sample leaf values for the random forest models (colored circles), alongside results for XGBoost (brown triangle) and Mokapot SVM (pink triangle). For each minimum sample leaf value, 13 different maximum depth settings were evaluated. The dashed red line indicates the 1% entrapment FDR threshold. Across all search engines and datasets, increasing model complexity generally results in a higher number of entrapment PSMs, particularly for highly complex random forest models and XGBoost.

PXD004565

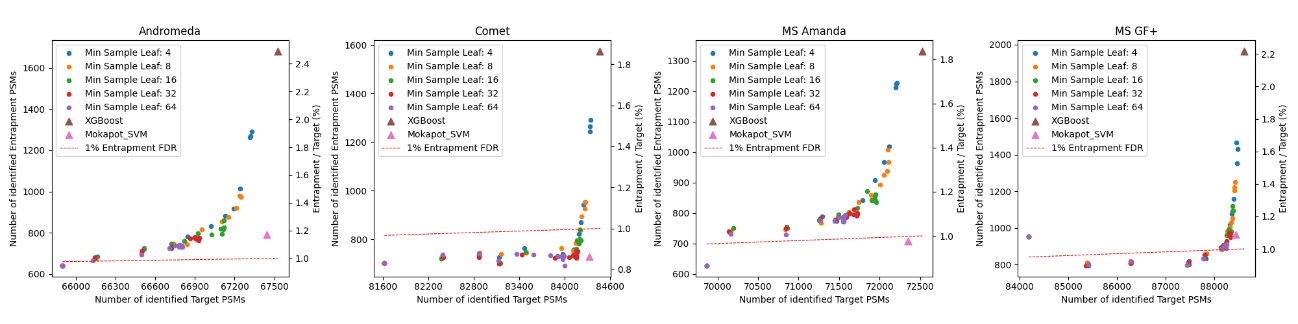

PXD004325

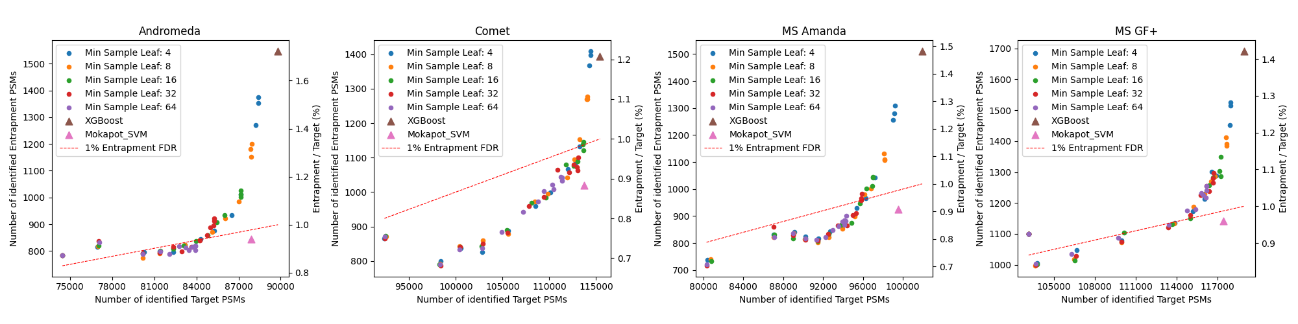

PXD009815

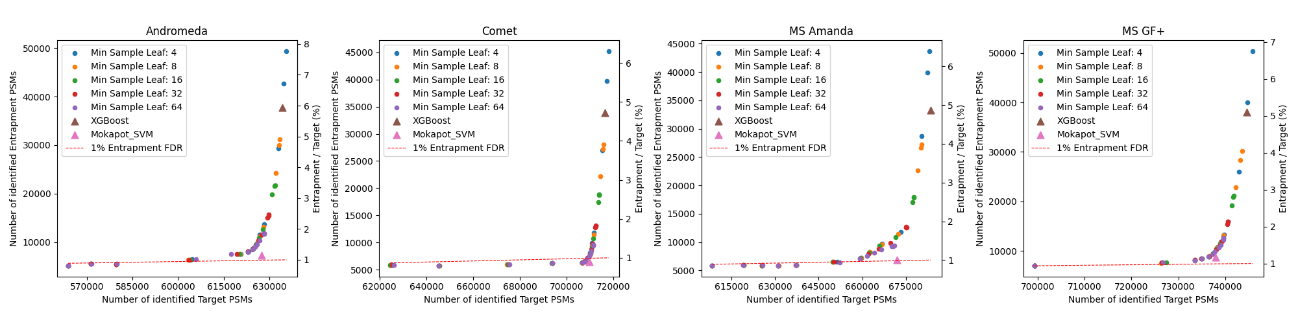

Supplementary Figure 5: Relationship between the number of identified target and entrapment PSMs for different model complexities when rescoring search results for datasets PXD004565, PXD004324, and PXD009815. Each panel corresponds to a different search engine (Andromeda, Comet, MS Amanda, and MS-GF+), displaying the distribution of identified PSMs for various rescoring models. Each point represents a model with different minimum sample leaf values for the random forest models (colored circles), alongside results for XGBoost (brown triangle) and Mokapot SVM (pink triangle). For each minimum sample leaf value, 13 different maximum depth settings were evaluated. The dashed red line indicates the 1% entrapment FDR threshold. Across all search engines and datasets, increasing model complexity generally results in a higher number of entrapment PSMs, particularly for highly complex random forest models and XGBoost.

PXD000561

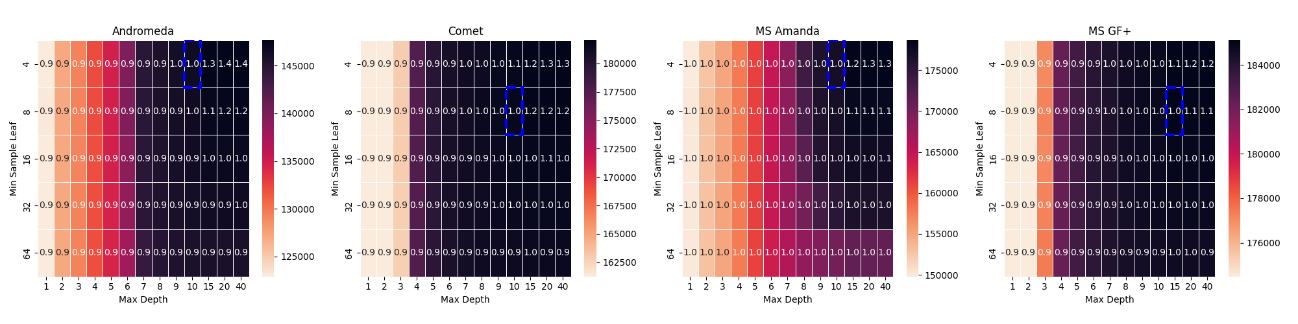

PXD006675

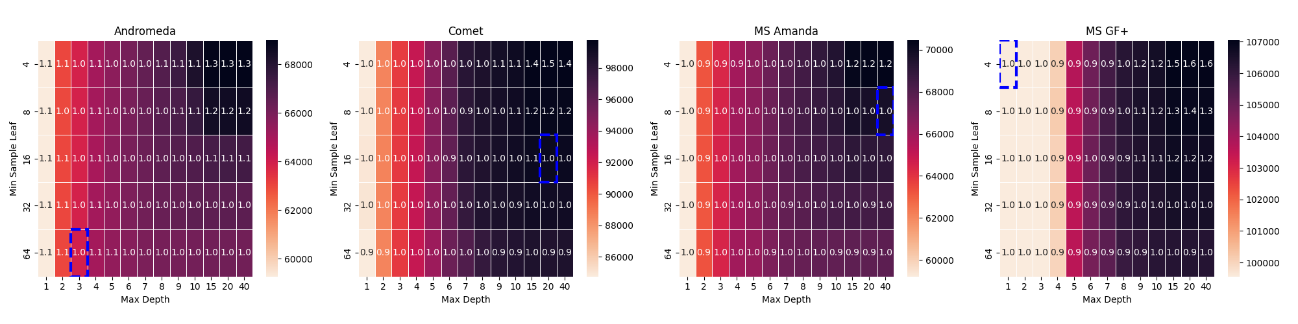

PXD001468

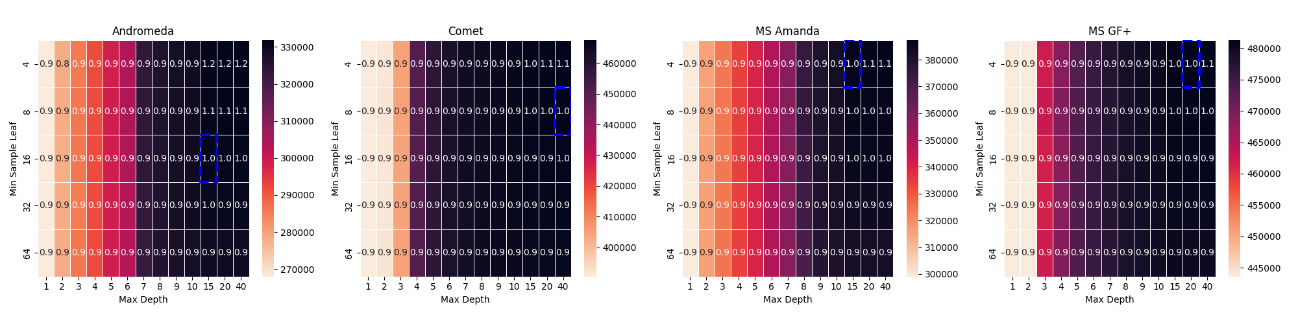

PXD001250

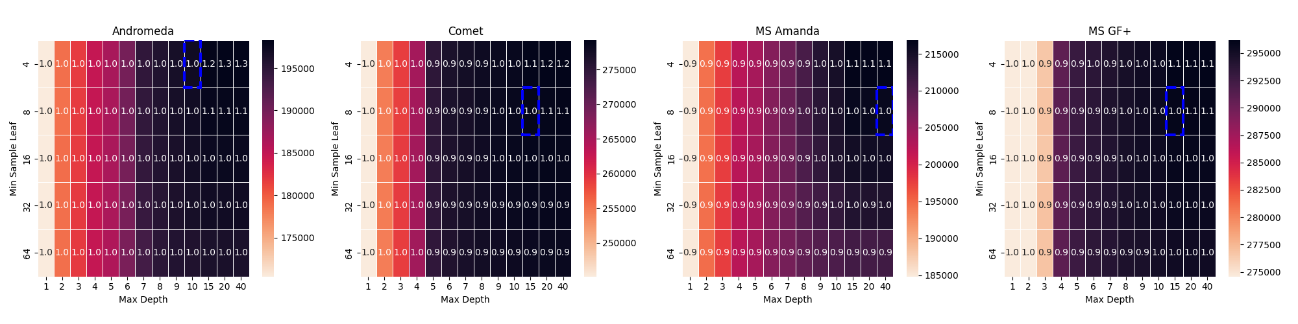

Supplementary Figure 6: Heatmaps showing the distribution of identified target PSMs across different hyperparameter settings for the random forest model when rescoring search results for datasets PXD000561, PXD006675, PXD001468, and PXD001250. The x-axis represents the maximum tree depth, while the y-axis corresponds to the minimum sample leaf parameter. Lighter colors indicate a higher number of identified target PSMs, with each cell annotated by the corresponding entrapment FDR value. The dashed blue rectangle highlights the optimal hyperparameter region, where the highest number of target PSMs is achieved while maintaining an entrapment FDR closest to 1%. Across all datasets, more complex models tend to exceed the expected entrapment FDR threshold, particularly at lower minimum sample leaf values and higher tree depths.

PXD000612

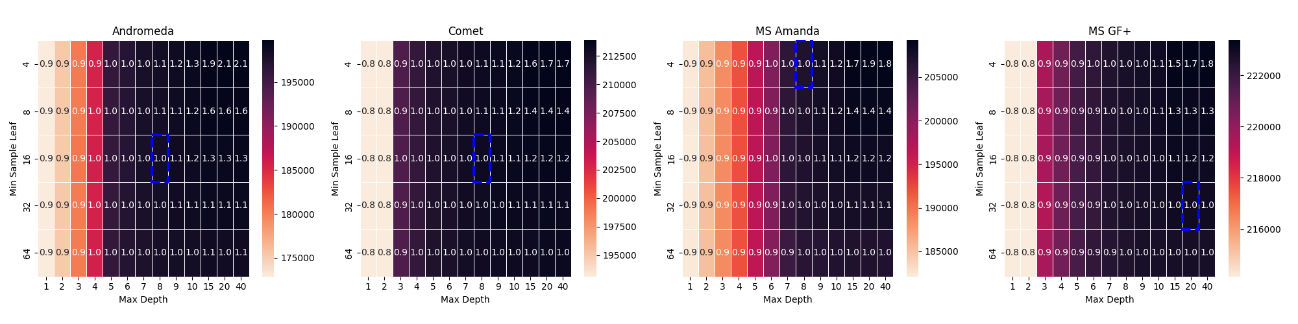

PXD004948

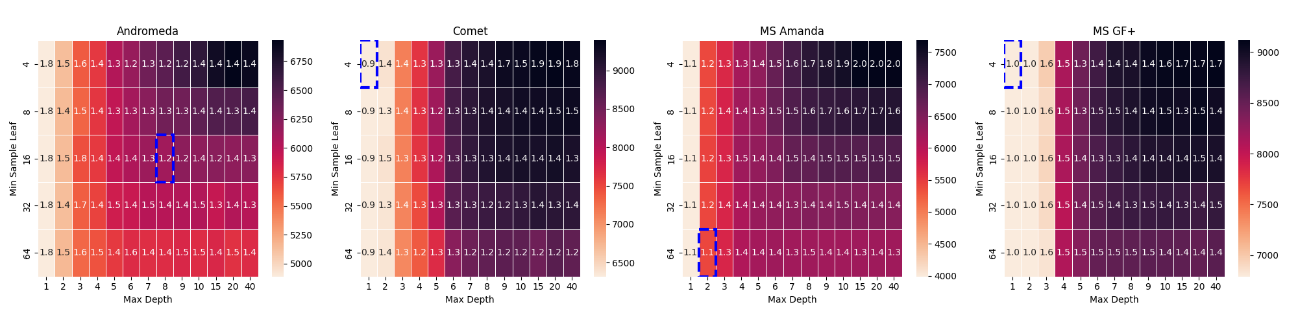

PXD040344

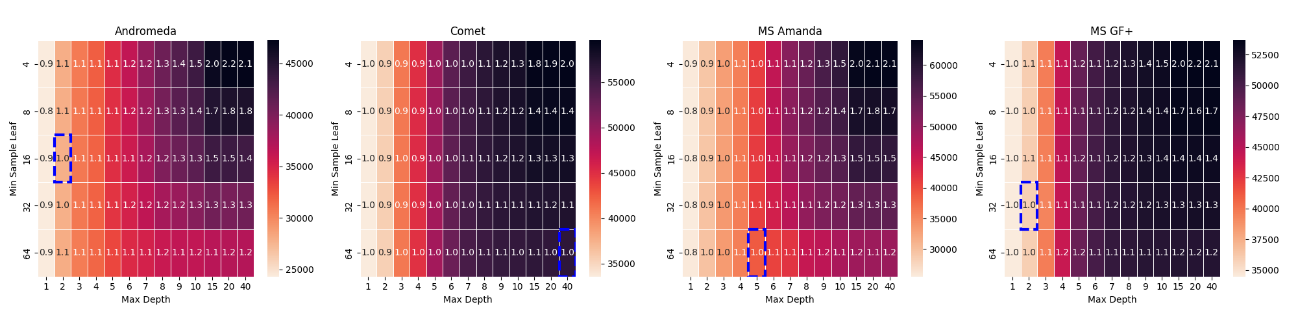

PXD004947

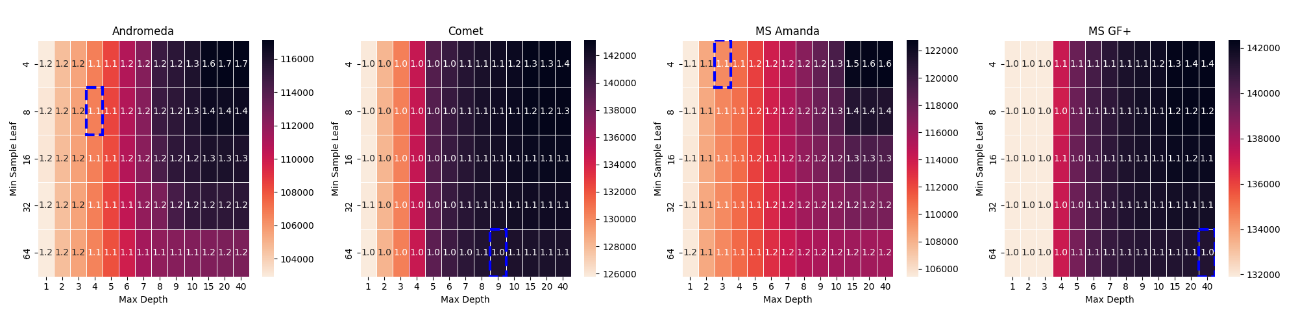

Supplementary Figure 7: Heatmaps showing the distribution of identified target PSMs across different hyperparameter settings for the random forest model when rescoring search results for PXD000612, PXD004948, PXD040344, and PXD004947. The x-axis represents the maximum tree depth, while the y-axis corresponds to the minimum sample leaf parameter. Lighter colors indicate a higher number of identified target PSMs, with each cell annotated by the corresponding entrapment FDR value. The dashed blue rectangle highlights the optimal hyperparameter region, where the highest number of target PSMs is achieved while maintaining an entrapment FDR closest to 1%. Across all datasets, more complex models tend to exceed the expected entrapment FDR threshold, particularly at lower minimum sample leaf values and higher tree depths.

PXD004565

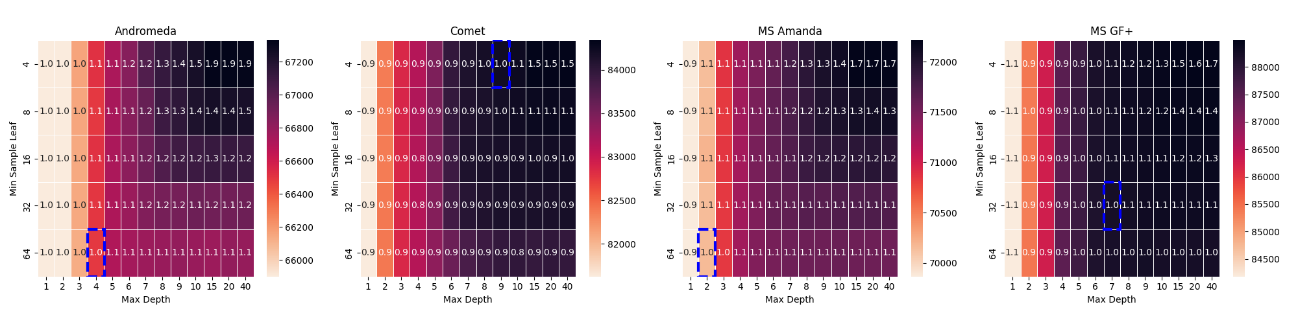

PXD004325

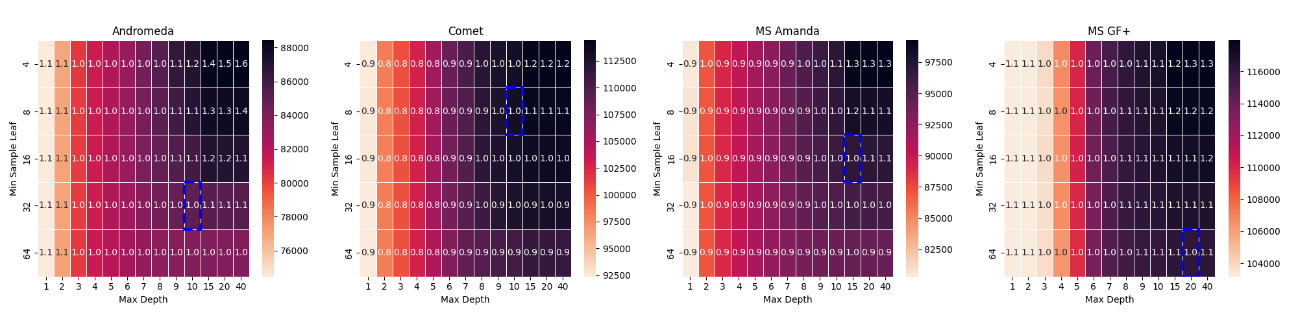

PXD009815

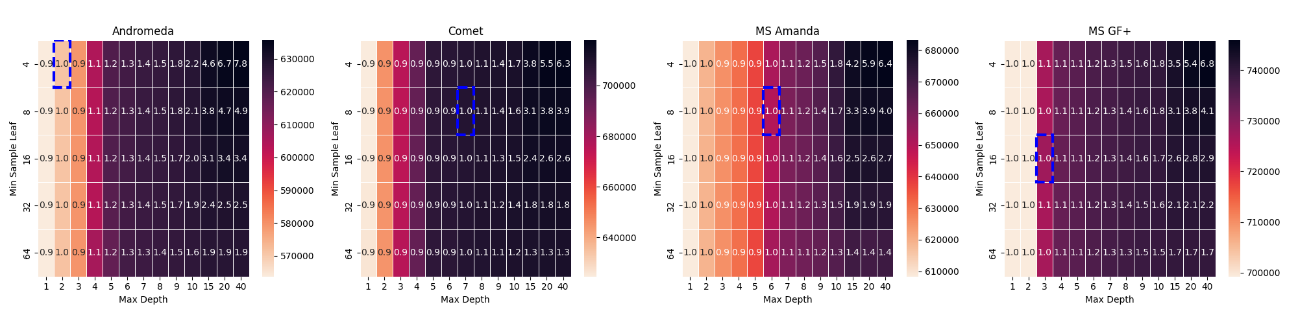

Supplementary Figure 8: Heatmaps showing the distribution of identified target PSMs across different hyperparameter settings for the random forest model when rescoring search results for PXD004565, PXD004324, and PXD009815. The x-axis represents the maximum tree depth, while the y-axis corresponds to the minimum sample leaf parameter. Lighter colors indicate a higher number of identified target PSMs, with each cell annotated by the corresponding entrapment FDR value. The dashed blue rectangle highlights the optimal hyperparameter region, where the highest number of target PSMs is achieved while maintaining an entrapment FDR closest to 1%. Across all datasets, more complex models tend to exceed the expected entrapment FDR threshold, particularly at lower minimum sample leaf values and higher tree depths.

PXD000561

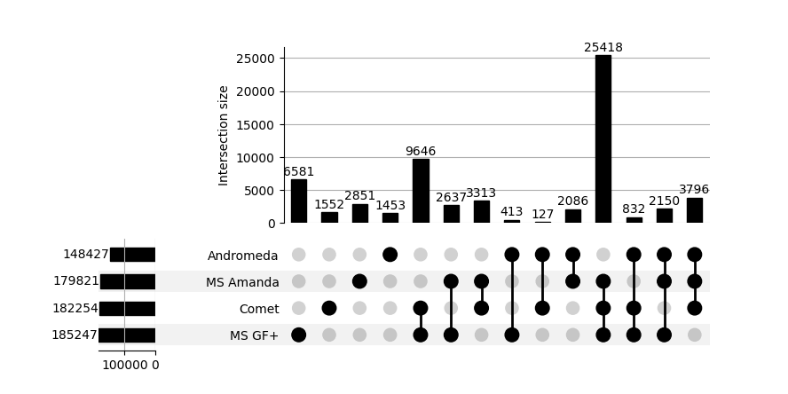

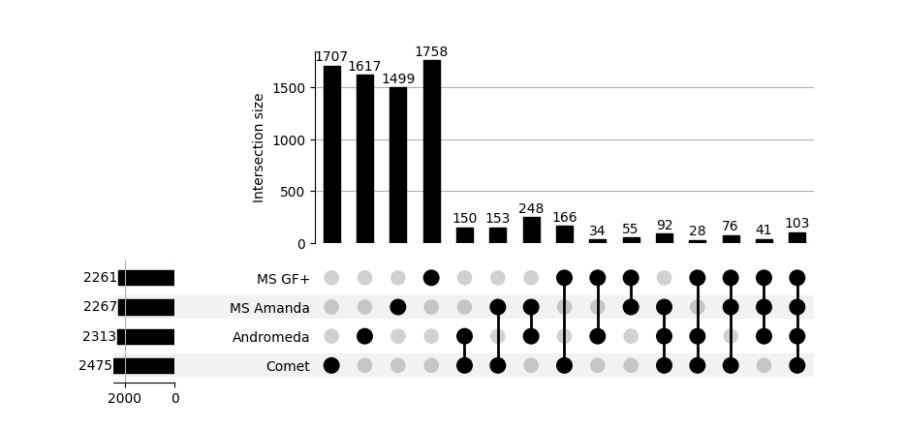

PXD006675

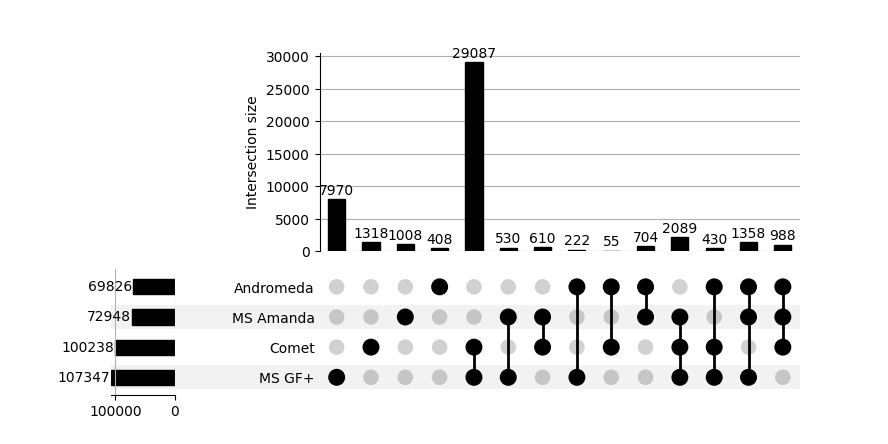

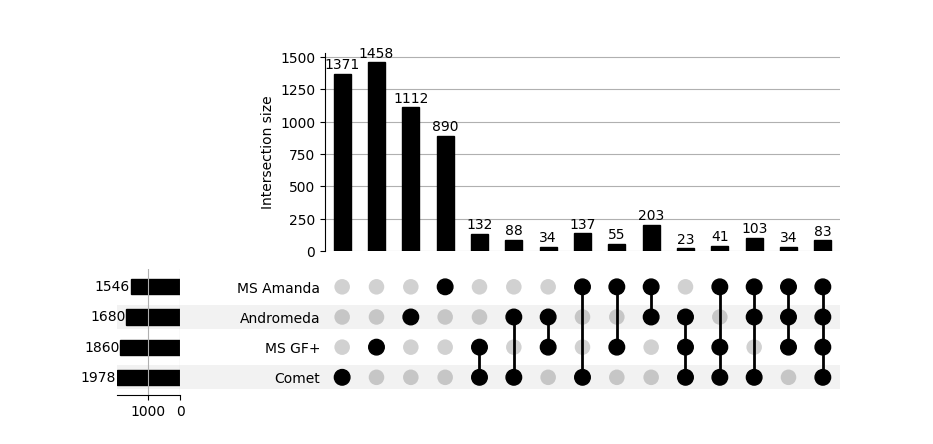

Supplementary Figure 9: UpSet plots illustrating the intersection of identified PSMs across different search engines after rescoring with XGBoost across datasets PXD000561, and PXD006675. The top panel represents target PSMs, while the down panel shows entrapment PSMs. The x-axis of each plot displays unique and shared intersections among the search engines, while the y-axis indicates the intersection size. The results highlight a strong overlap among target PSM identifications across search engines, whereas entrapment PSMs appear mostly unique to each search engine, suggesting that most entrapment matches are randomly distributed. For the target PSMs figure, the intersection of all search engines, containing the largest number of PSMs, was excluded to improve visualization and facilitate comparison of smaller intersections.

PXD001468

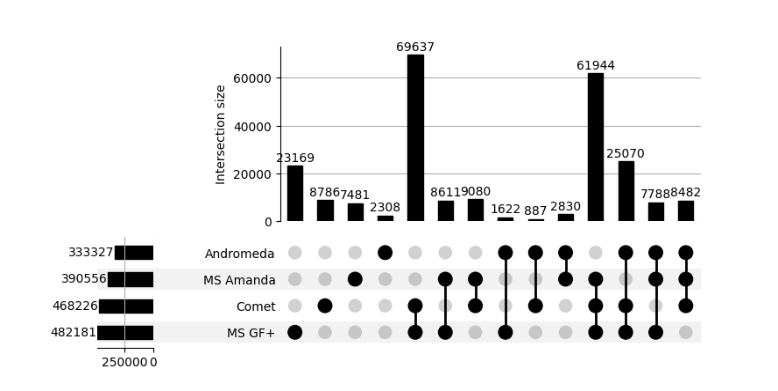

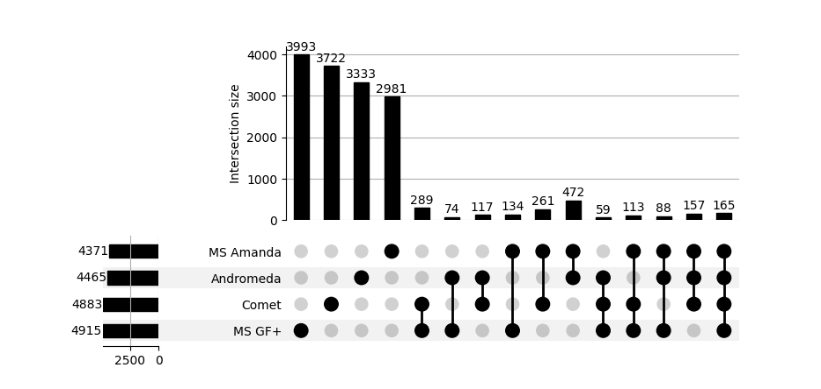

PXD001250

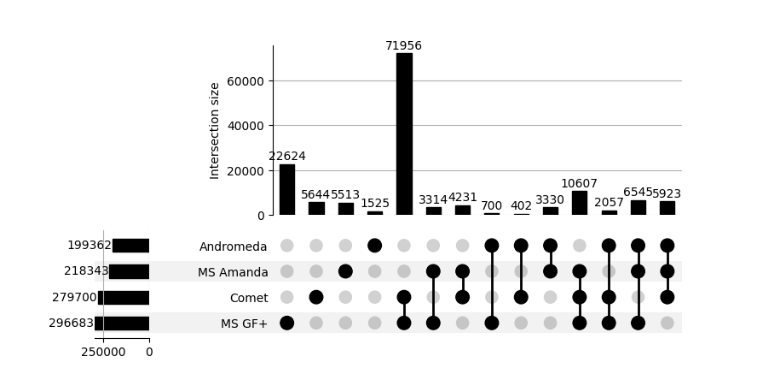

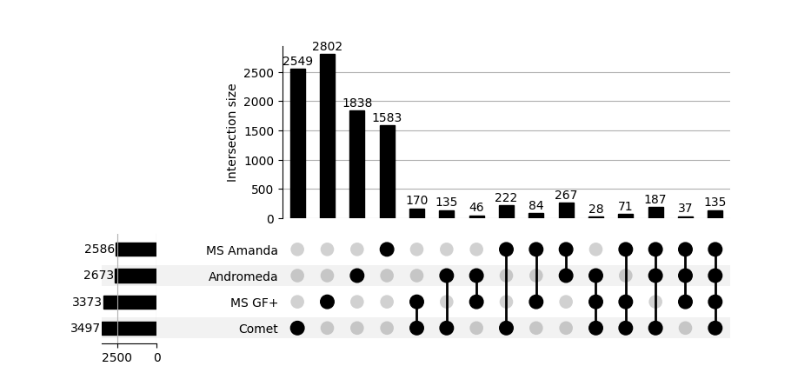

Supplementary Figure 10: UpSet plots illustrating the intersection of identified PSMs across different search engines after rescoring with XGBoost across datasets PXD001468, and PXD001250. The top panel represents target PSMs, while the down panel shows entrapment PSMs. The x-axis of each plot displays unique and shared intersections among the search engines, while the y-axis indicates the intersection size. The results highlight a strong overlap among target PSM identifications across search engines, whereas entrapment PSMs appear mostly unique to each search engine, suggesting that most entrapment matches are randomly distributed. For the target PSMs figure, the intersection of all search engines, containing the largest number of PSMs, was excluded to improve visualization and facilitate comparison of smaller intersections.

PXD000612

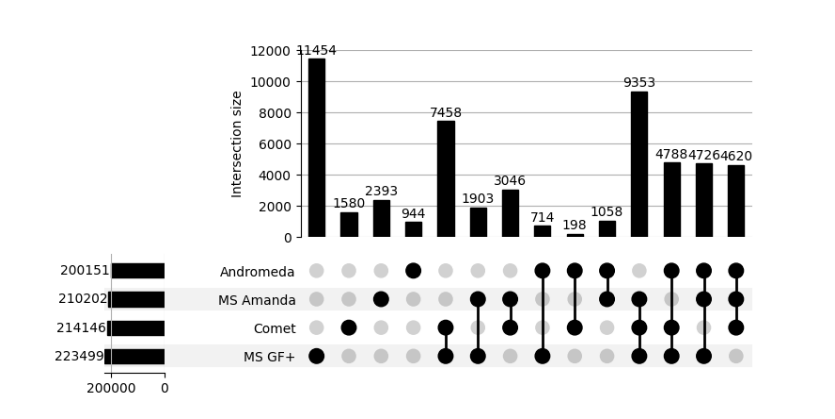

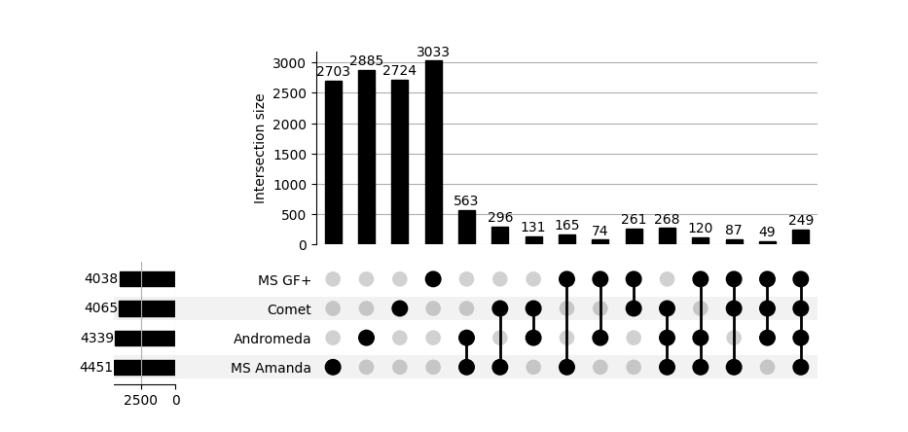

PXD004948

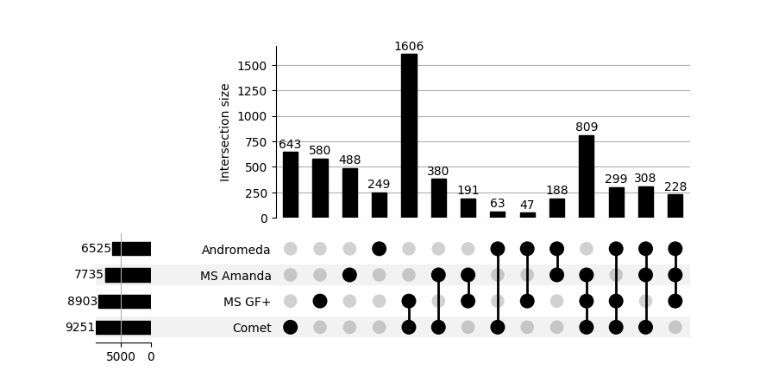

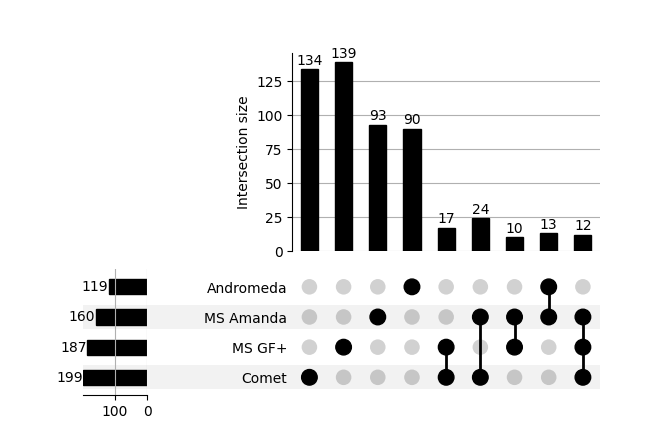

Supplementary Figure 11: UpSet plots illustrating the intersection of identified PSMs across different search engines after rescoring with XGBoost across datasets PXD000612, and PXD004948. The top panel represents target PSMs, while the down panel shows entrapment PSMs. The x-axis of each plot displays unique and shared intersections among the search engines, while the y-axis indicates the intersection size. The results highlight a strong overlap among target PSM identifications across search engines, whereas entrapment PSMs appear mostly unique to each search engine, suggesting that most entrapment matches are randomly distributed. For the target PSMs figure, the intersection of all search engines, containing the largest number of PSMs, was excluded to improve visualization and facilitate comparison of smaller intersections.

PXD040344

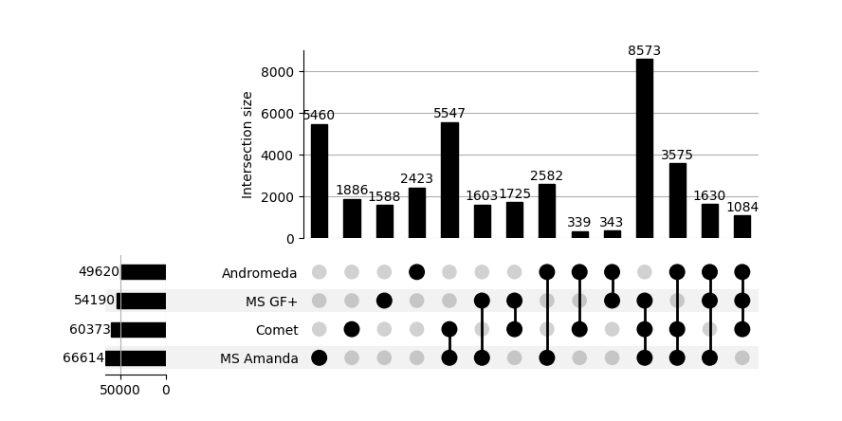

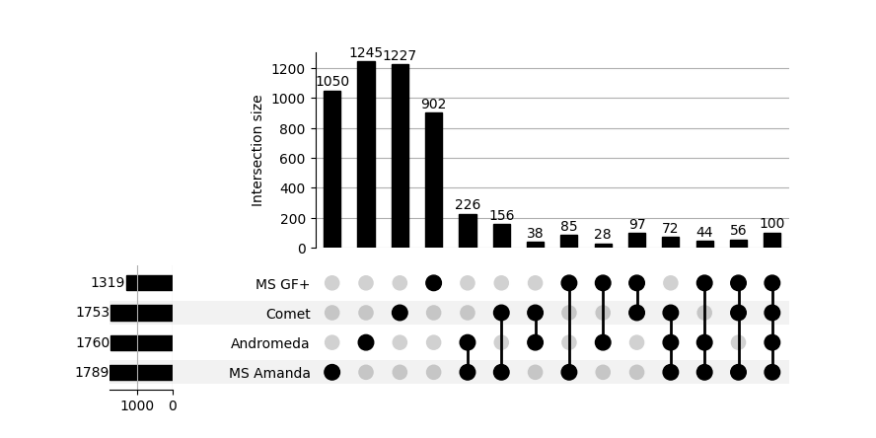

PXD004947

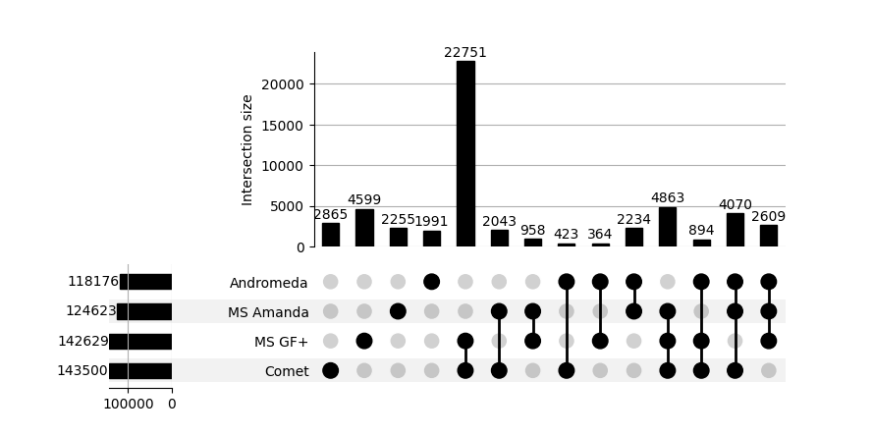

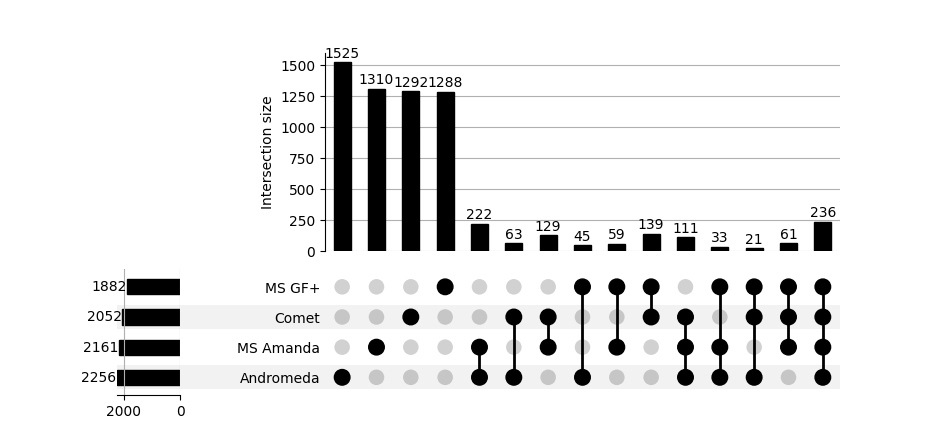

Supplementary Figure 12: UpSet plots illustrating the intersection of identified PSMs across different search engines after rescoring with XGBoost across datasets PXD040344, and PXD004947. The top panel represents target PSMs, while the down panel shows entrapment PSMs. The x-axis of each plot displays unique and shared intersections among the search engines, while the y-axis indicates the intersection size. The results highlight a strong overlap among target PSM identifications across search engines, whereas entrapment PSMs appear mostly unique to each search engine, suggesting that most entrapment matches are randomly distributed. For the target PSMs figure, the intersection of all search engines, containing the largest number of PSMs, was excluded to improve visualization and facilitate comparison of smaller intersections.

PXD004565

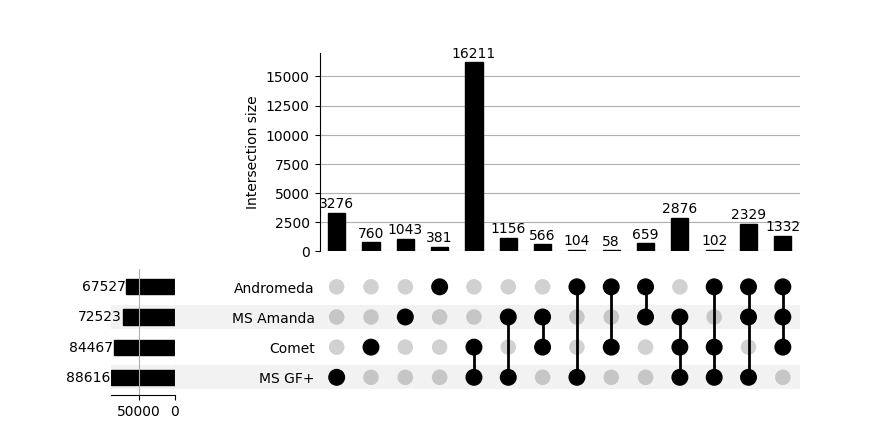

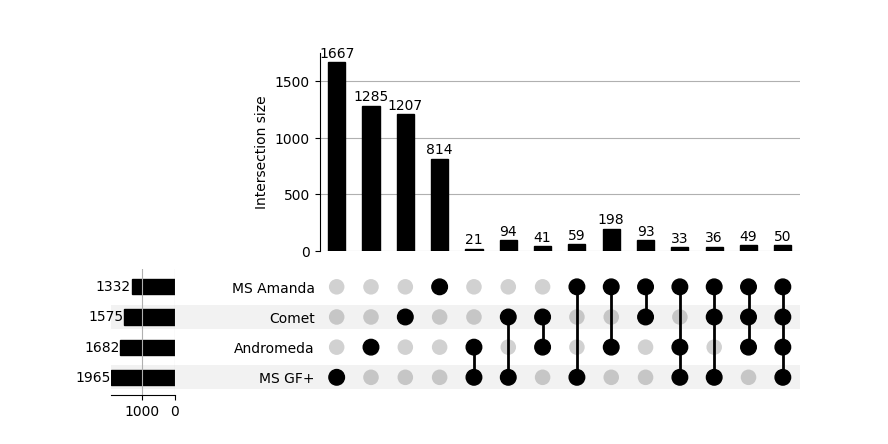

PXD004325

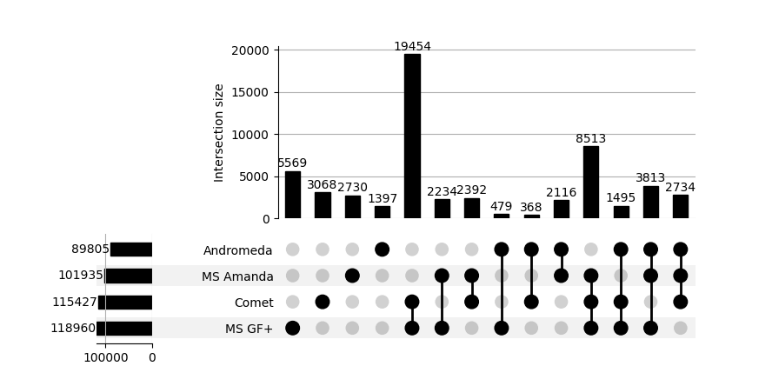

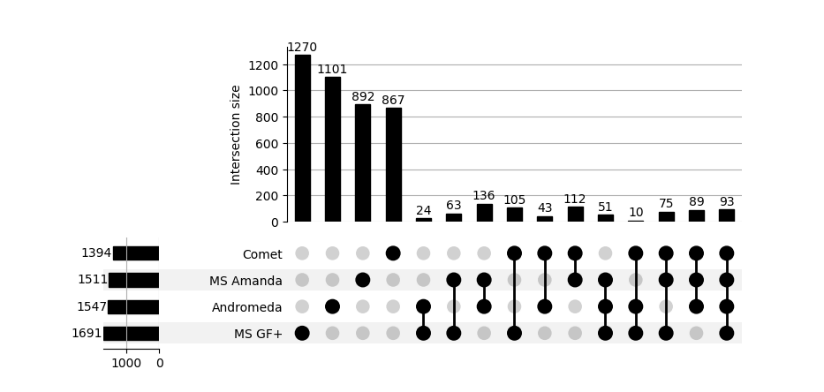

Supplementary Figure 13: UpSet plots illustrating the intersection of identified PSMs across different search engines after rescoring with XGBoost across datasets PXD004565, and PXD004325. The top panel represents target PSMs, while the down panel shows entrapment PSMs. The x-axis of each plot displays unique and shared intersections among the search engines, while the y-axis indicates the intersection size. The results highlight a strong overlap among target PSM identifications across search engines, whereas entrapment PSMs appear mostly unique to each search engine, suggesting that most entrapment matches are randomly distributed. For the target PSMs figure, the intersection of all search engines, containing the largest number of PSMs, was excluded to improve visualization and facilitate comparison of smaller intersections.

PXD009815

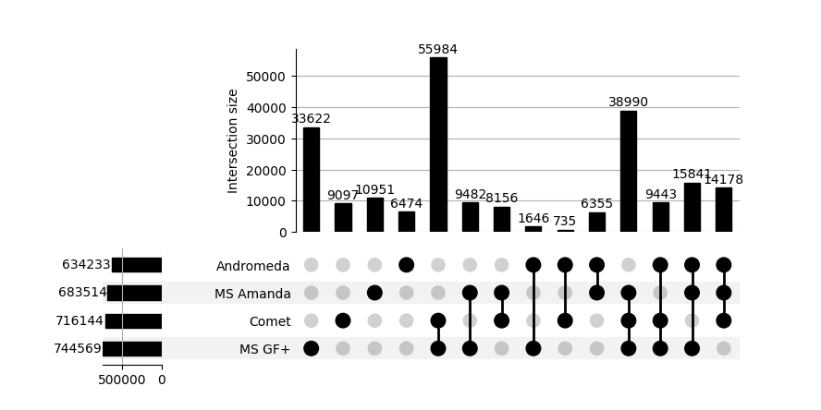

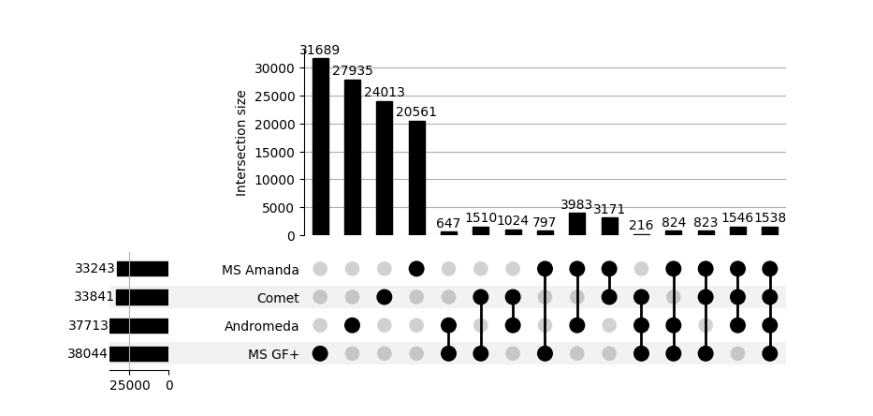

Supplementary Figure 14: UpSet plots illustrating the intersection of identified PSMs across different search engines after rescoring with XGBoost across dataset PXD009815. The top panel represents target PSMs, while the down panel shows entrapment PSMs. The x-axis of each plot displays unique and shared intersections among the search engines, while the y-axis indicates the intersection size. The results highlight a strong overlap among target PSM identifications across search engines, whereas entrapment PSMs appear mostly unique to each search engine, suggesting that most entrapment matches are randomly distributed. For the target PSMs figure, the intersection of all search engines, containing the largest number of PSMs, was excluded to improve visualization and facilitate comparison of smaller intersections.

PXD000561

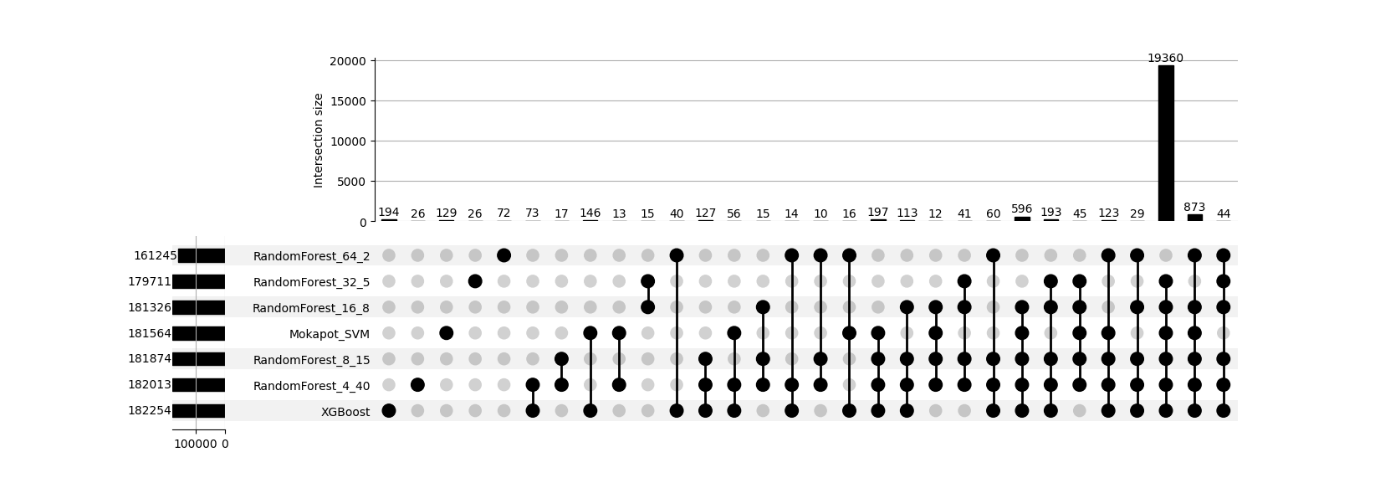

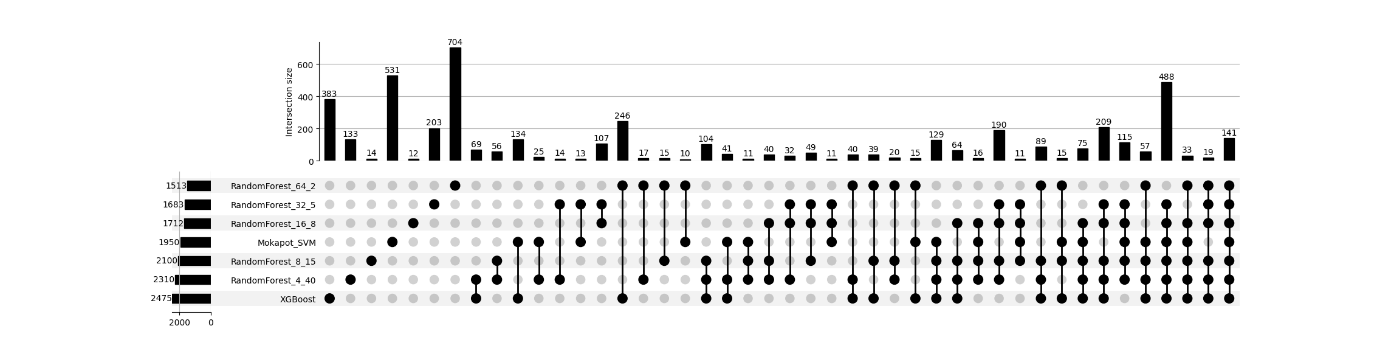

PXD006675

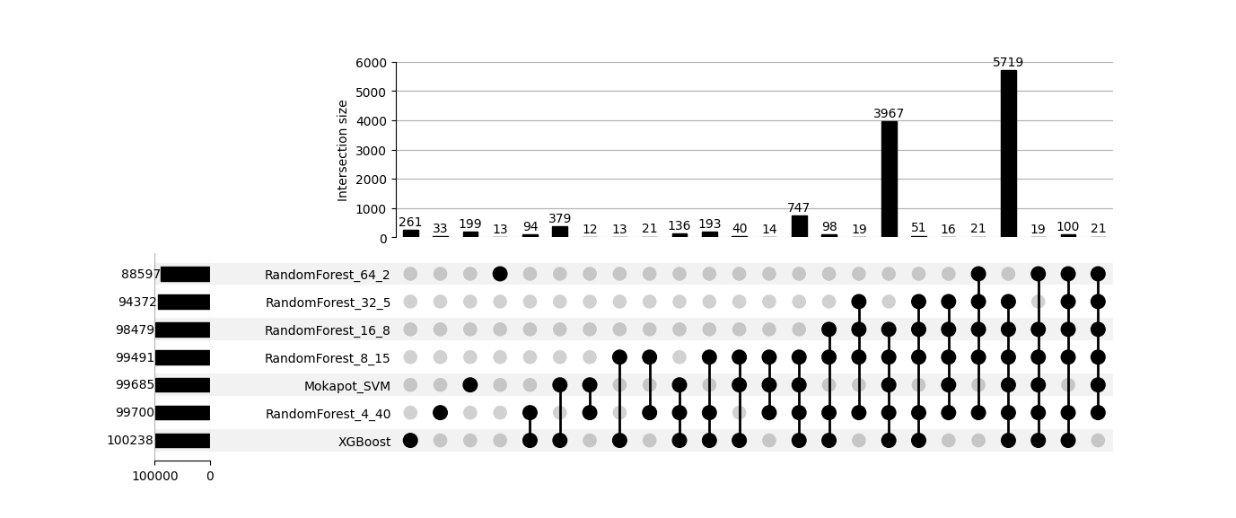

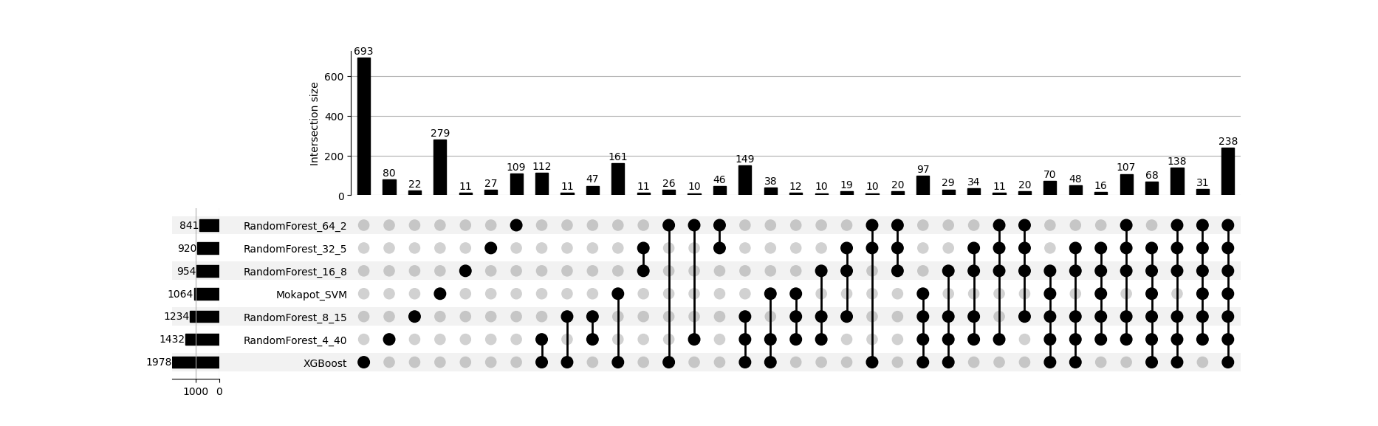

Supplementary Figure 15: UpSet plots illustrating the intersection of identified PSMs across different rescoring algorithms for Comet search results across datasets PXD000561, and PXD006675. The top panel represents target PSMs, while the down panel shows entrapment PSMs. The x-axis of each plot displays unique and shared intersections among the search engines, while the y-axis indicates the intersection size. The results highlight a strong overlap among target PSM identifications across search engines, whereas entrapment PSMs appear mostly unique to each search engine, suggesting that most entrapment matches are randomly distributed. For the target PSMs figure, the intersection of all search engines, containing the largest number of PSMs, was excluded to improve visualization and facilitate comparison of smaller intersections.

PXD001468

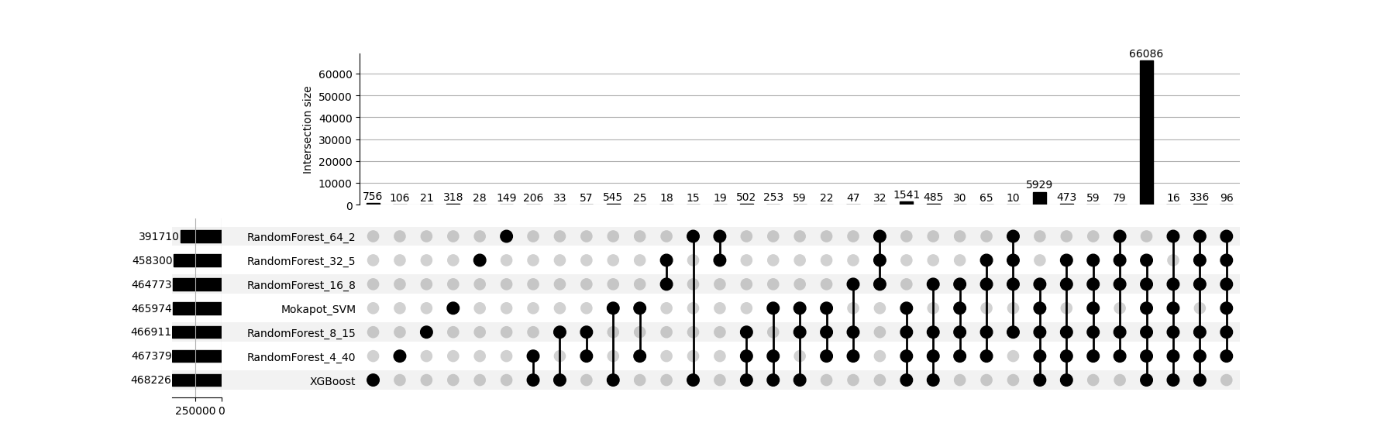

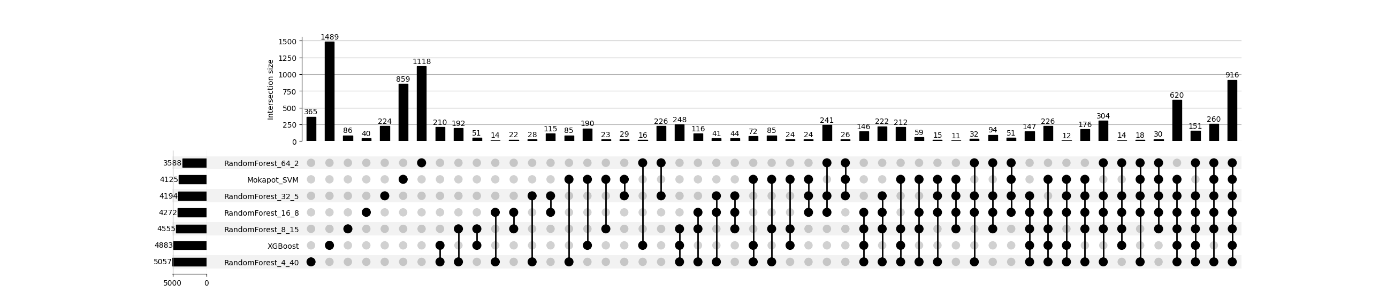

PXD001250

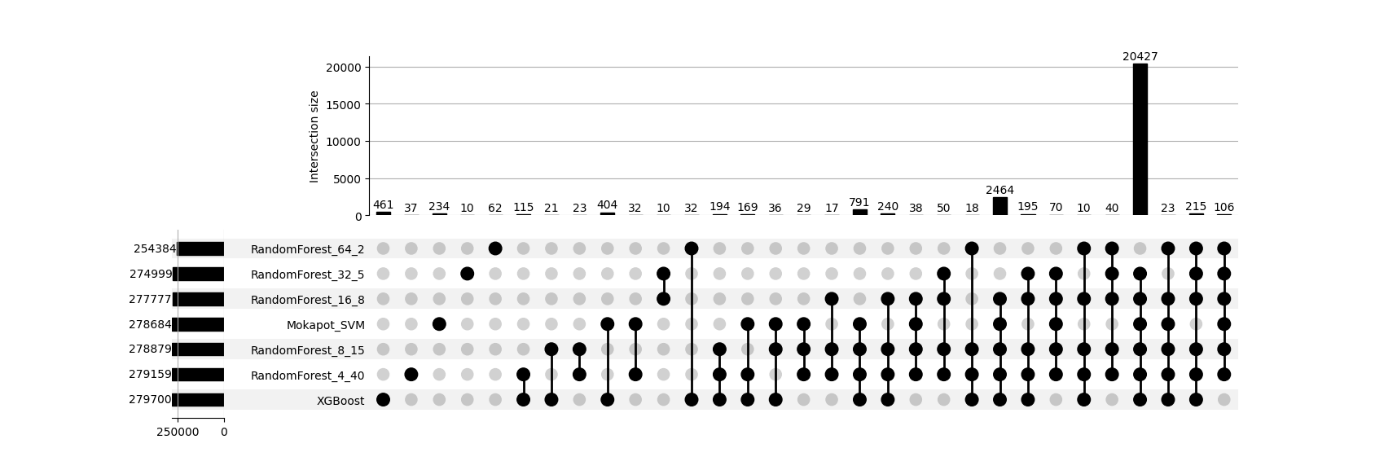

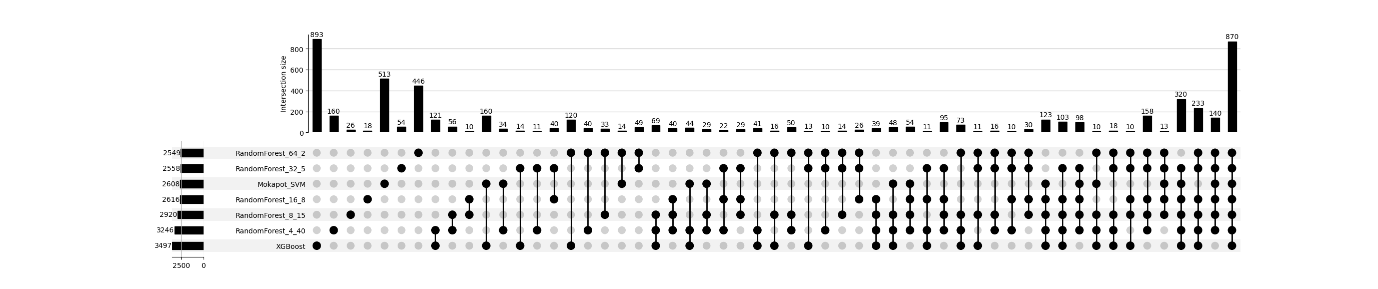

Supplementary Figure 16: UpSet plots illustrating the intersection of identified PSMs across different rescoring algorithms for Comet search results across datasets PXD001468, and PXD001250. The top panel represents target PSMs, while the down panel shows entrapment PSMs. The x-axis of each plot displays unique and shared intersections among the search engines, while the y-axis indicates the intersection size. The results highlight a strong overlap among target PSM identifications across search engines, whereas entrapment PSMs appear mostly unique to each search engine, suggesting that most entrapment matches are randomly distributed. For the target PSMs figure, the intersection of all search engines, containing the largest number of PSMs, was excluded to improve visualization and facilitate comparison of smaller intersections.

PXD000612

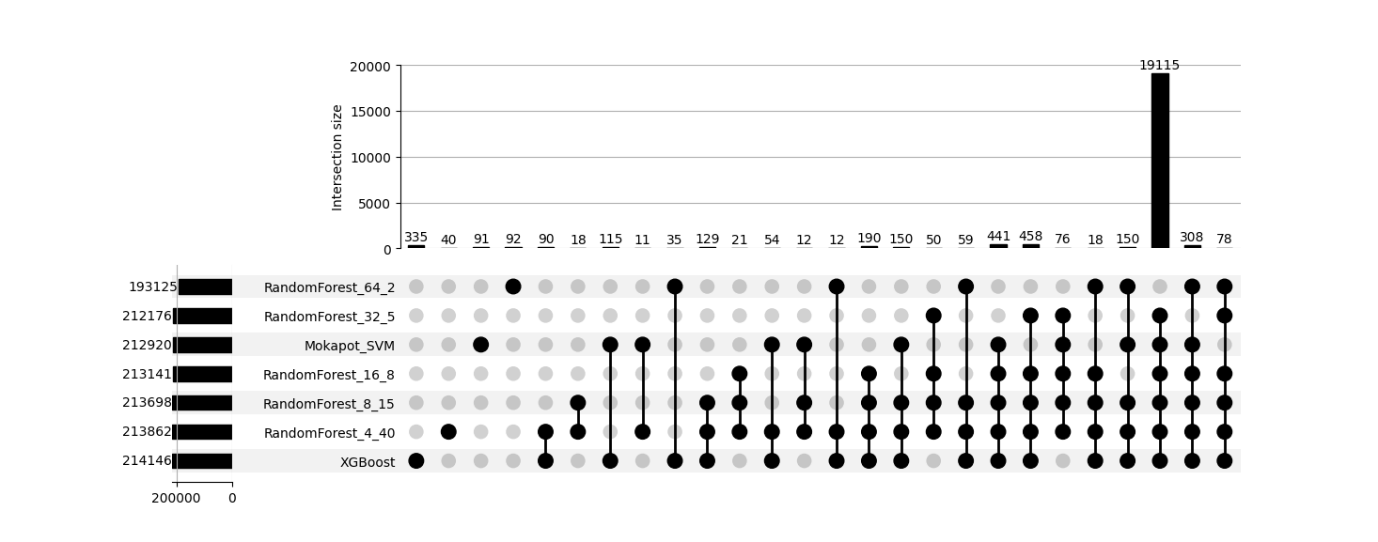

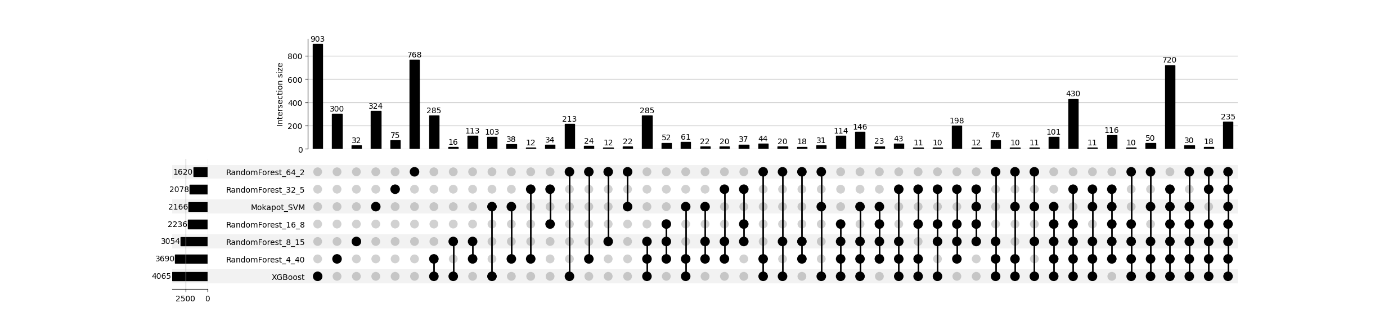

PXD004948

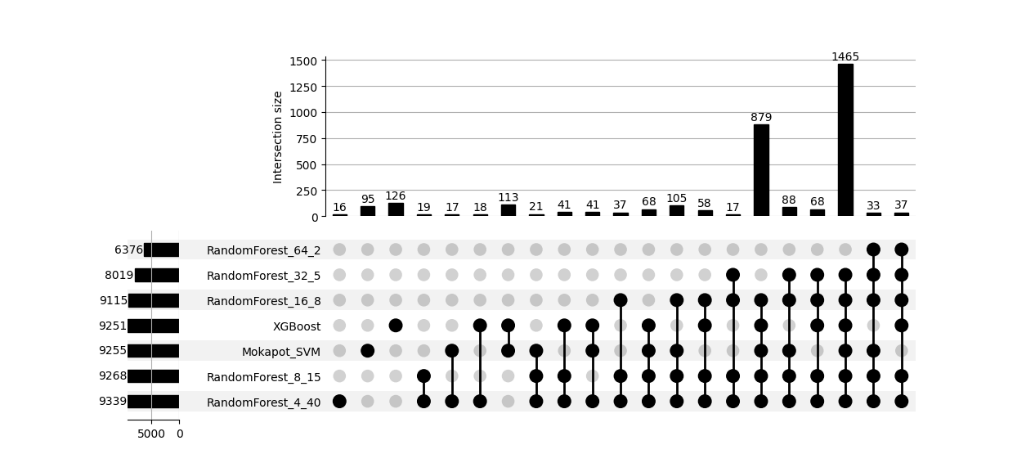

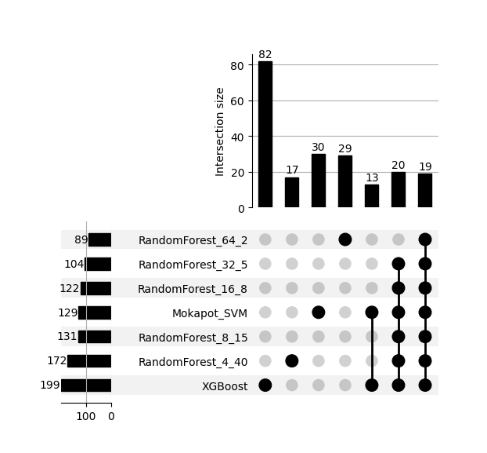

Supplementary Figure 17: UpSet plots illustrating the intersection of identified PSMs across different rescoring algorithms for Comet search results across datasets PXD000612, and PXD004948. The top panel represents target PSMs, while the down panel shows entrapment PSMs. The x-axis of each plot displays unique and shared intersections among the search engines, while the y-axis indicates the intersection size. The results highlight a strong overlap among target PSM identifications across search engines, whereas entrapment PSMs appear mostly unique to each search engine, suggesting that most entrapment matches are randomly distributed. For the target PSMs figure, the intersection of all search engines, containing the largest number of PSMs, was excluded to improve visualization and facilitate comparison of smaller intersections.

PXD040344

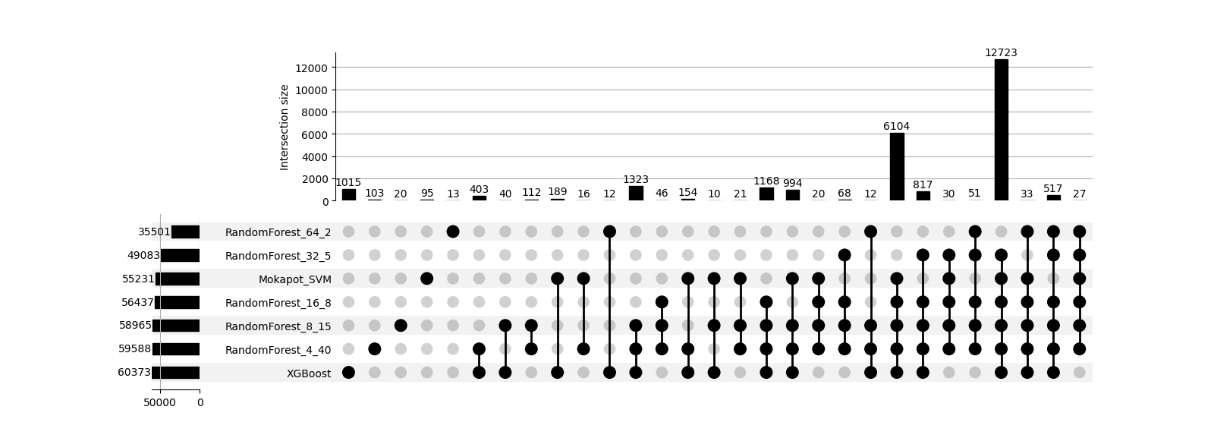

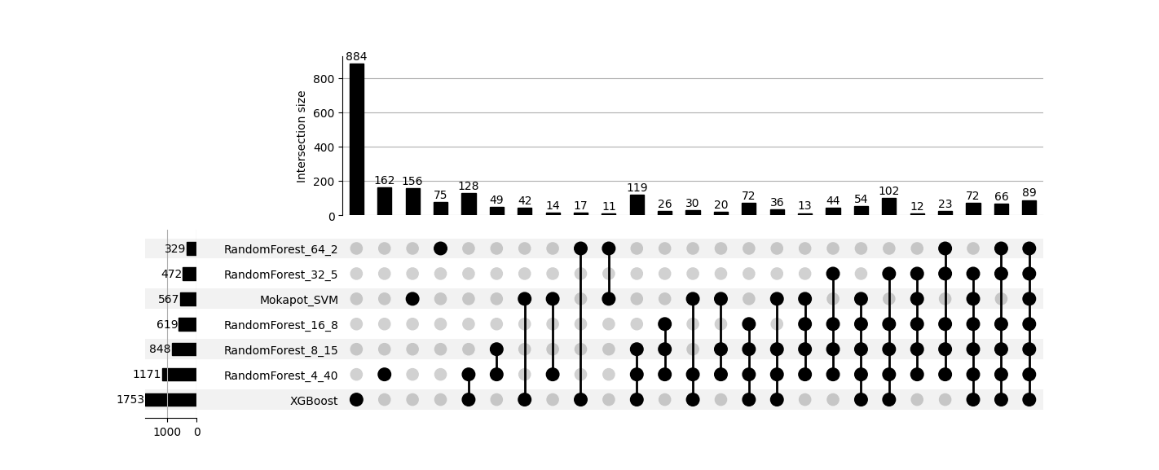

PXD004947

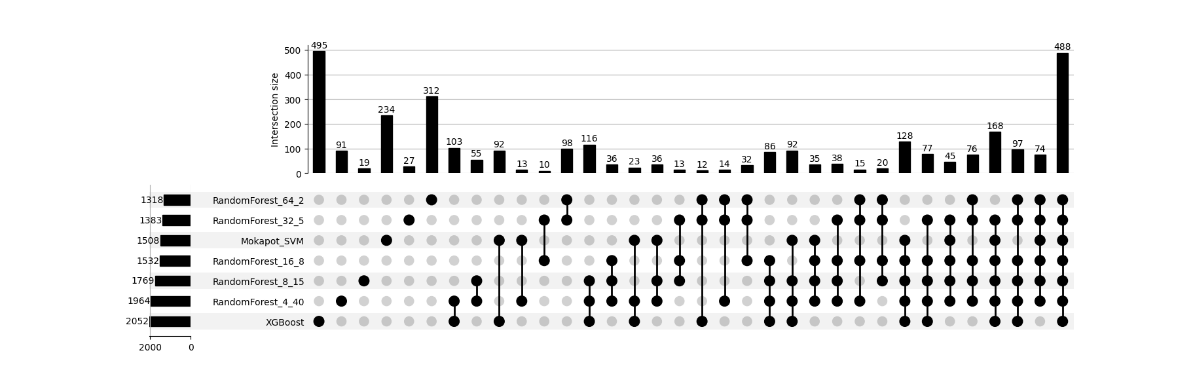

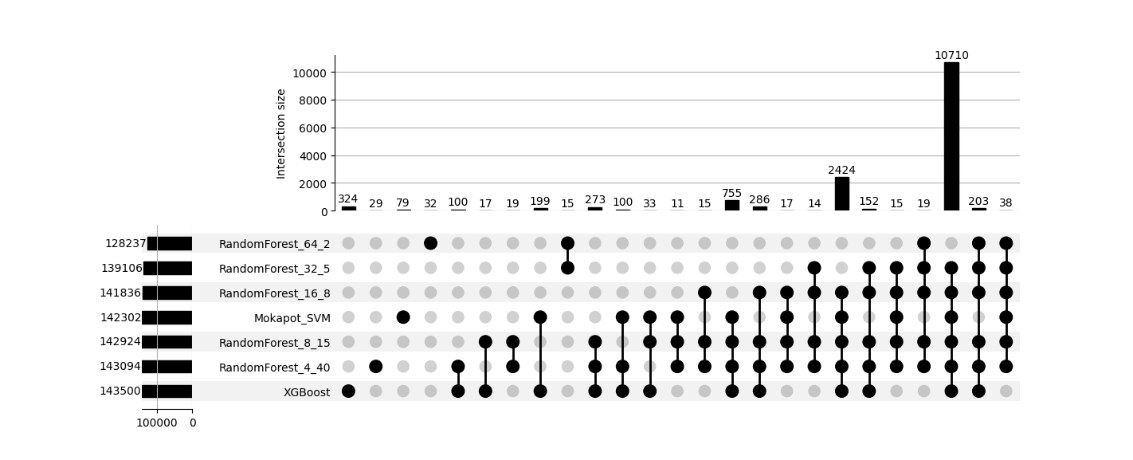

Supplementary Figure 18: UpSet plots illustrating the intersection of identified PSMs across different rescoring algorithms for Comet search results across datasets PXD040344, and PXD004947. The top panel represents target PSMs, while the down panel shows entrapment PSMs. The x-axis of each plot displays unique and shared intersections among the search engines, while the y-axis indicates the intersection size. The results highlight a strong overlap among target PSM identifications across search engines, whereas entrapment PSMs appear mostly unique to each search engine, suggesting that most entrapment matches are randomly distributed. For the target PSMs figure, the intersection of all search engines, containing the largest number of PSMs, was excluded to improve visualization and facilitate comparison of smaller intersections.

PXD004565

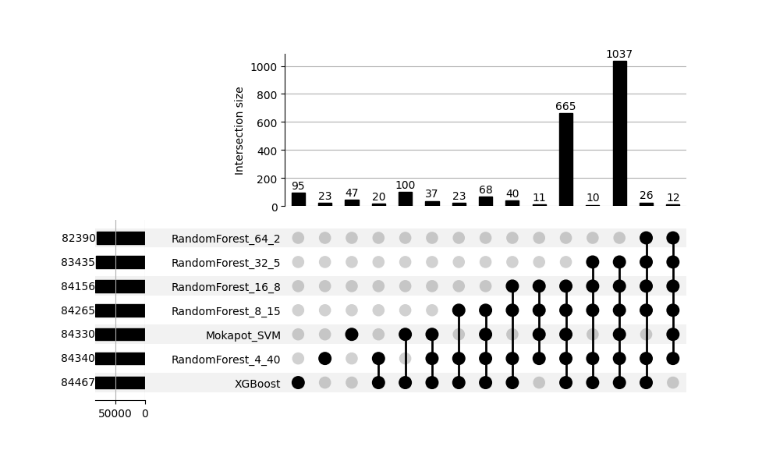

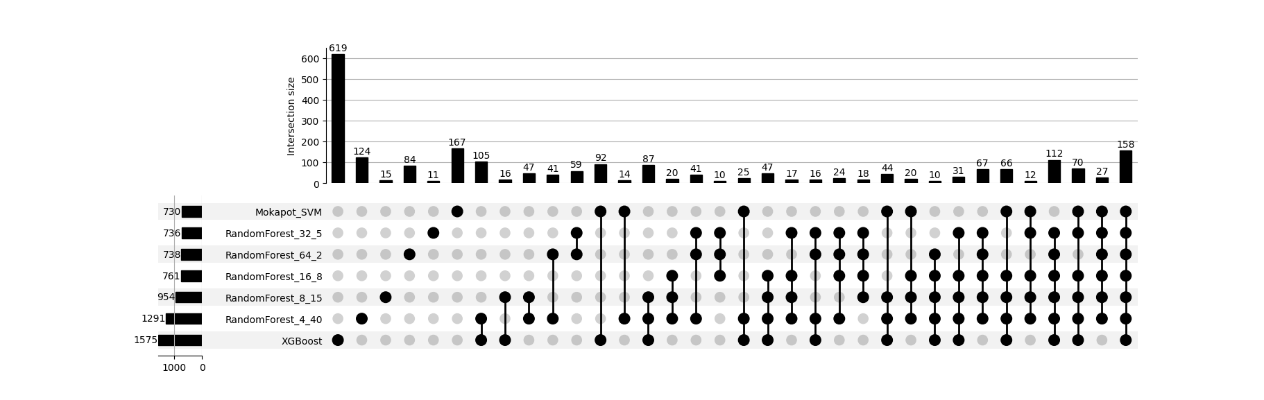

PXD004325

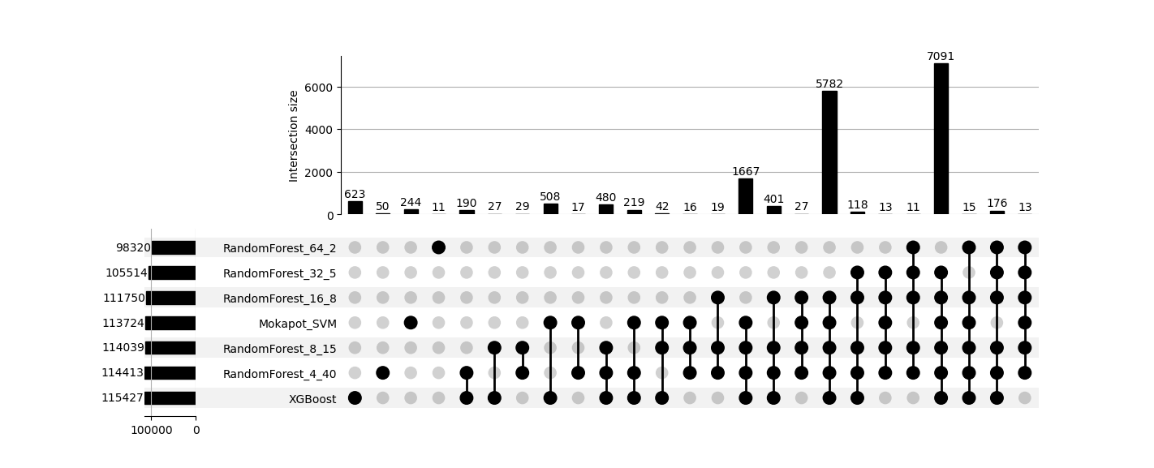

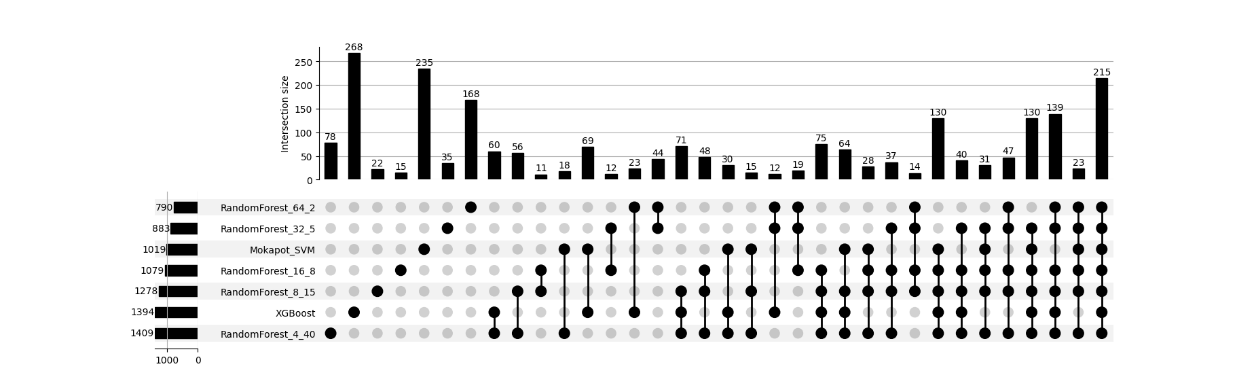

Supplementary Figure 19: UpSet plots illustrating the intersection of identified PSMs across different rescoring algorithms for Comet search results across datasets PXD004565, and PXD004325. The top panel represents target PSMs, while the down panel shows entrapment PSMs. The x-axis of each plot displays unique and shared intersections among the search engines, while the y-axis indicates the intersection size. The results highlight a strong overlap among target PSM identifications across search engines, whereas entrapment PSMs appear mostly unique to each search engine, suggesting that most entrapment matches are randomly distributed. For the target PSMs figure, the intersection of all search engines, containing the largest number of PSMs, was excluded to improve visualization and facilitate comparison of smaller intersections.

PXD009815

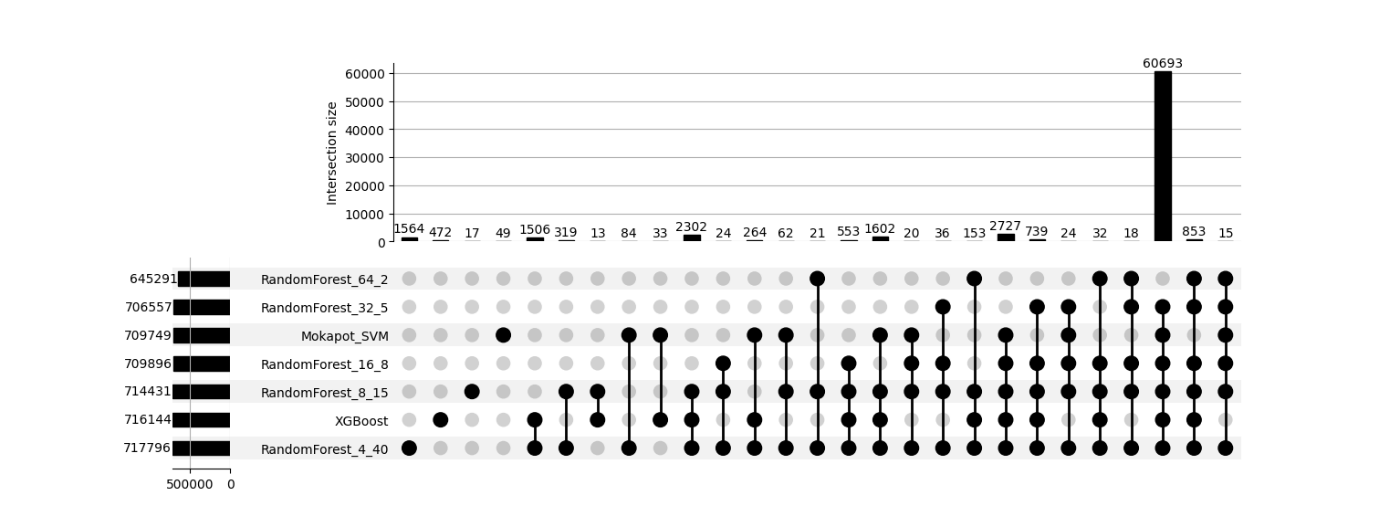

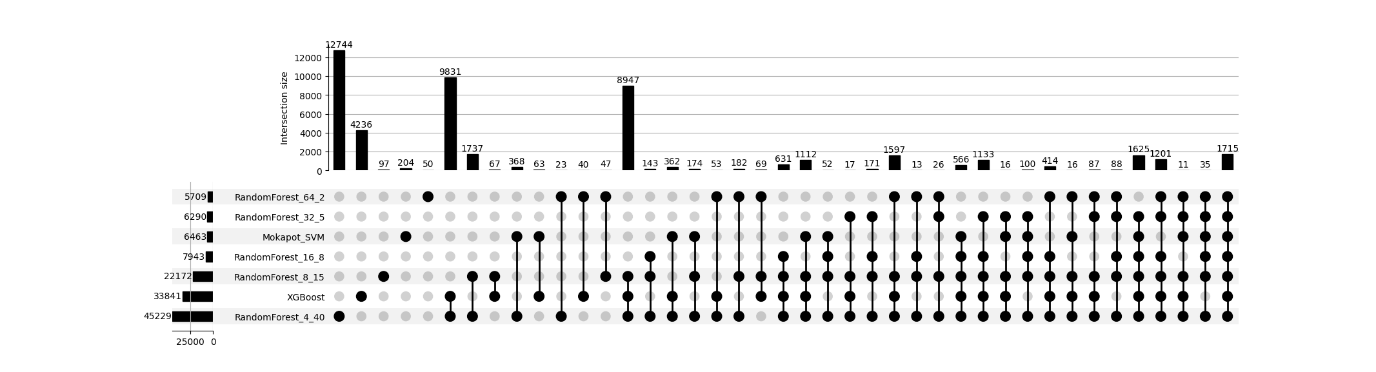

Supplementary Figure 20: UpSet plots illustrating the intersection of identified PSMs across different rescoring algorithms for Comet search results across dataset PXD009815. The top panel represents target PSMs, while the down panel shows entrapment PSMs. The x-axis of each plot displays unique and shared intersections among the search engines, while the y-axis indicates the intersection size. The results highlight a strong overlap among target PSM identifications across search engines, whereas entrapment PSMs appear mostly unique to each search engine, suggesting that most entrapment matches are randomly distributed. For the target PSMs figure, the intersection of all search engines, containing the largest number of PSMs, was excluded to improve visualization and facilitate comparison of smaller intersections.

PXD000561

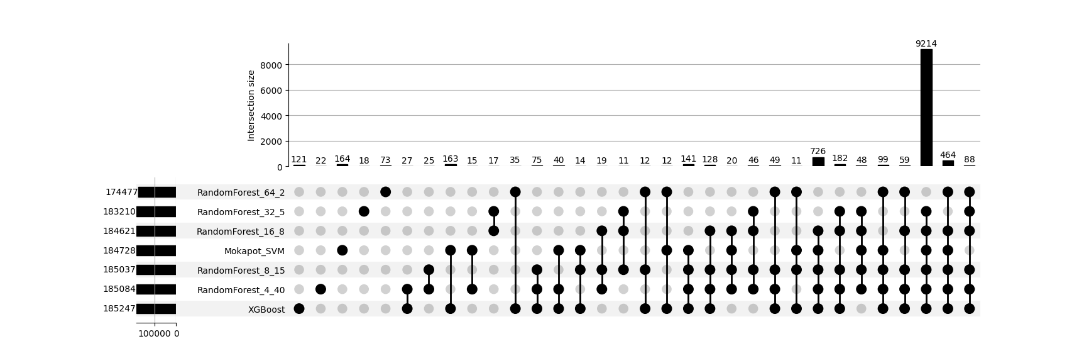

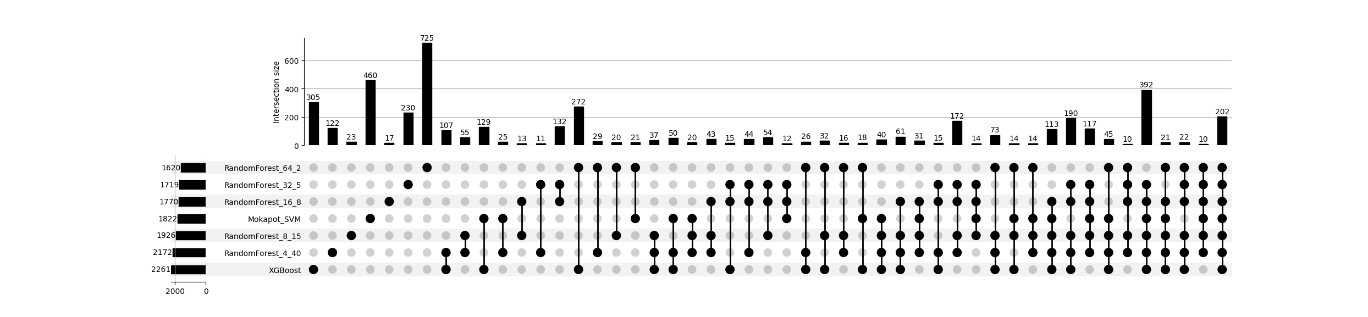

PXD006675

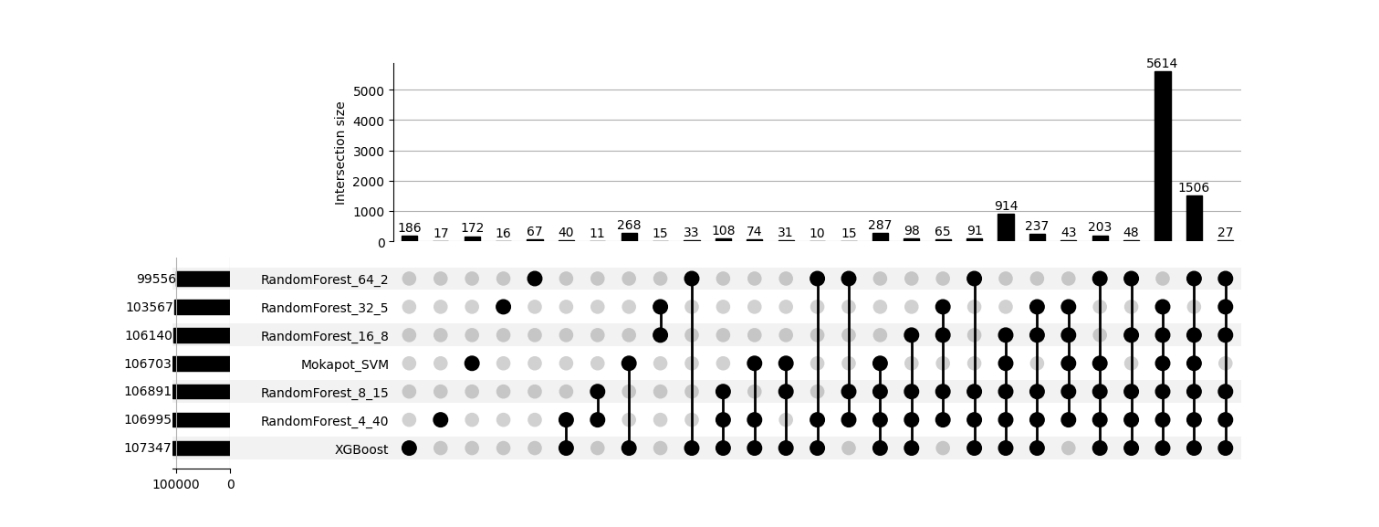

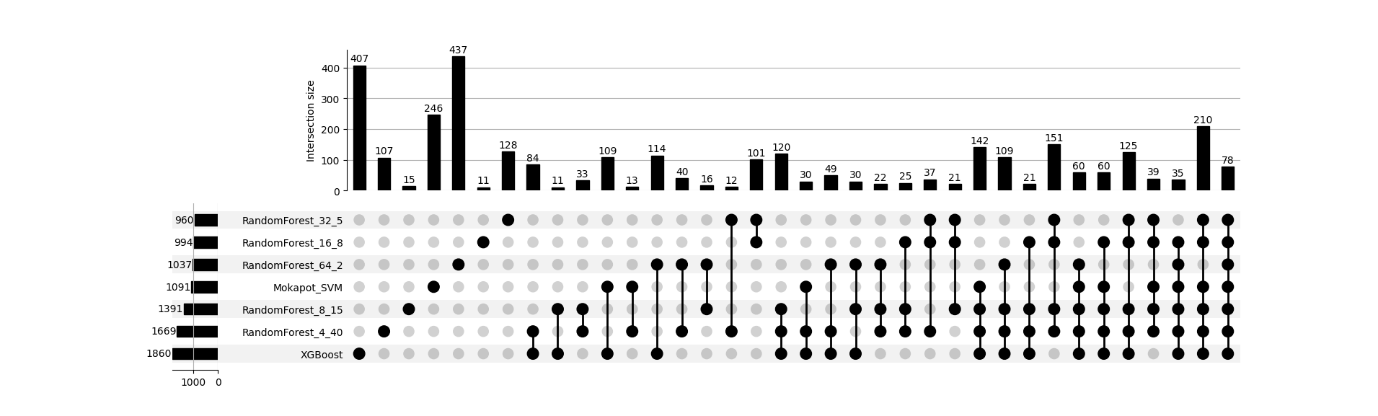

Supplementary Figure 21: UpSet plots illustrating the intersection of identified PSMs across different rescoring algorithms for MS-GF+ search results across datasets PXD000561, and PXD006675. The top panel represents target PSMs, while the down panel shows entrapment PSMs. The x-axis of each plot displays unique and shared intersections among the search engines, while the y-axis indicates the intersection size. The results highlight a strong overlap among target PSM identifications across search engines, whereas entrapment PSMs appear mostly unique to each search engine, suggesting that most entrapment matches are randomly distributed. For the target PSMs figure, the intersection of all search engines, containing the largest number of PSMs, was excluded to improve visualization and facilitate comparison of smaller intersections.

PXD001468

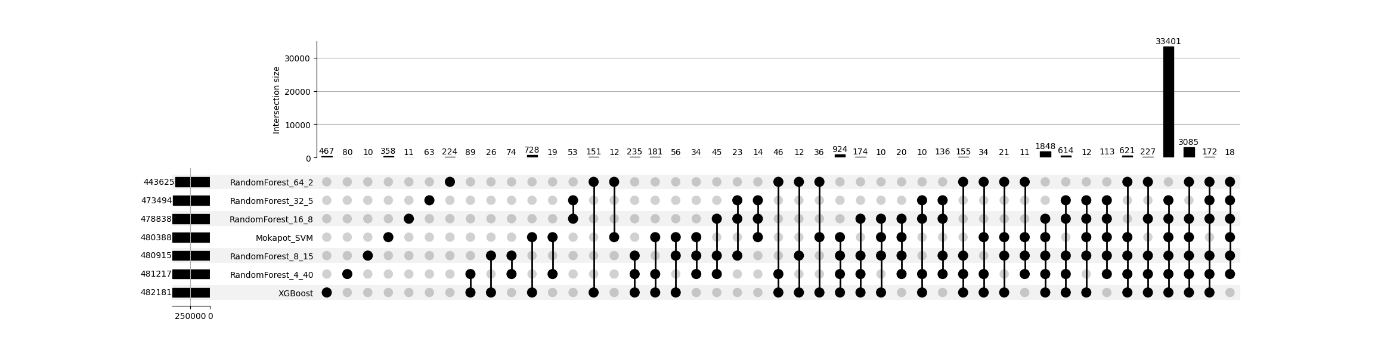

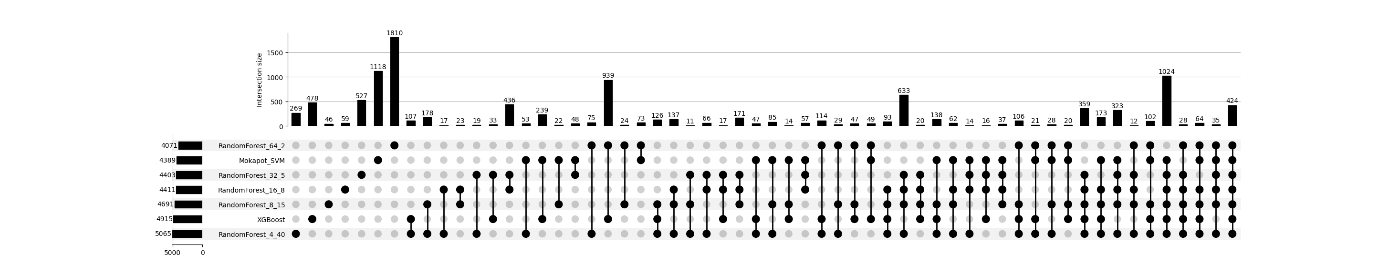

PXD001250

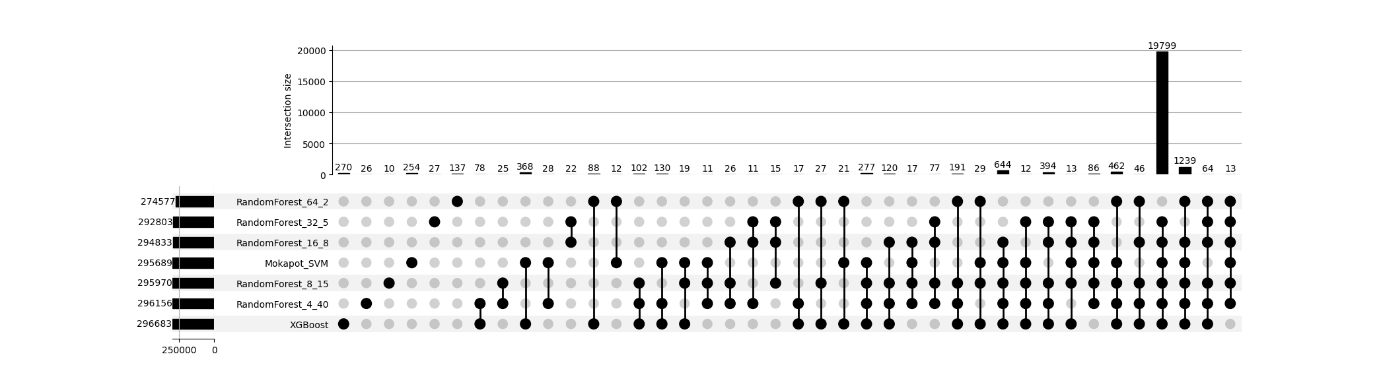

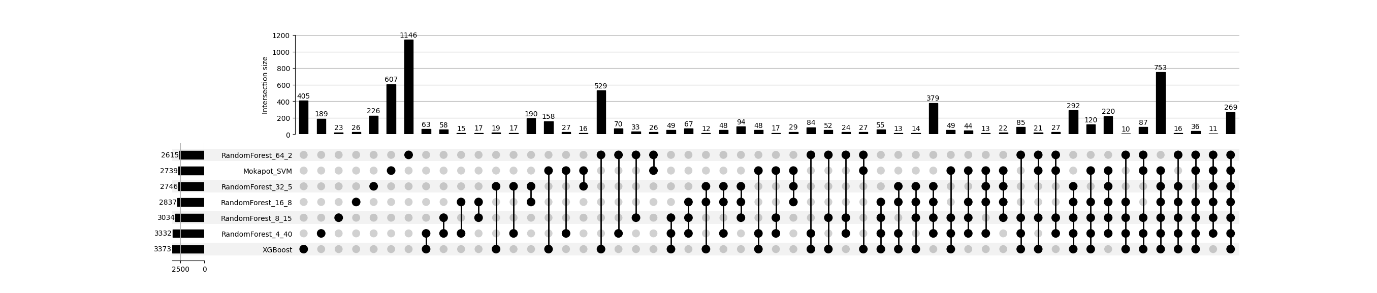

Supplementary Figure 22: UpSet plots illustrating the intersection of identified PSMs across different rescoring algorithms for MS-GF+ search results across datasets PXD001468, and PXD001250. The top panel represents target PSMs, while the down panel shows entrapment PSMs. The x-axis of each plot displays unique and shared intersections among the search engines, while the y-axis indicates the intersection size. The results highlight a strong overlap among target PSM identifications across search engines, whereas entrapment PSMs appear mostly unique to each search engine, suggesting that most entrapment matches are randomly distributed. For the target PSMs figure, the intersection of all search engines, containing the largest number of PSMs, was excluded to improve visualization and facilitate comparison of smaller intersections.

PXD000612

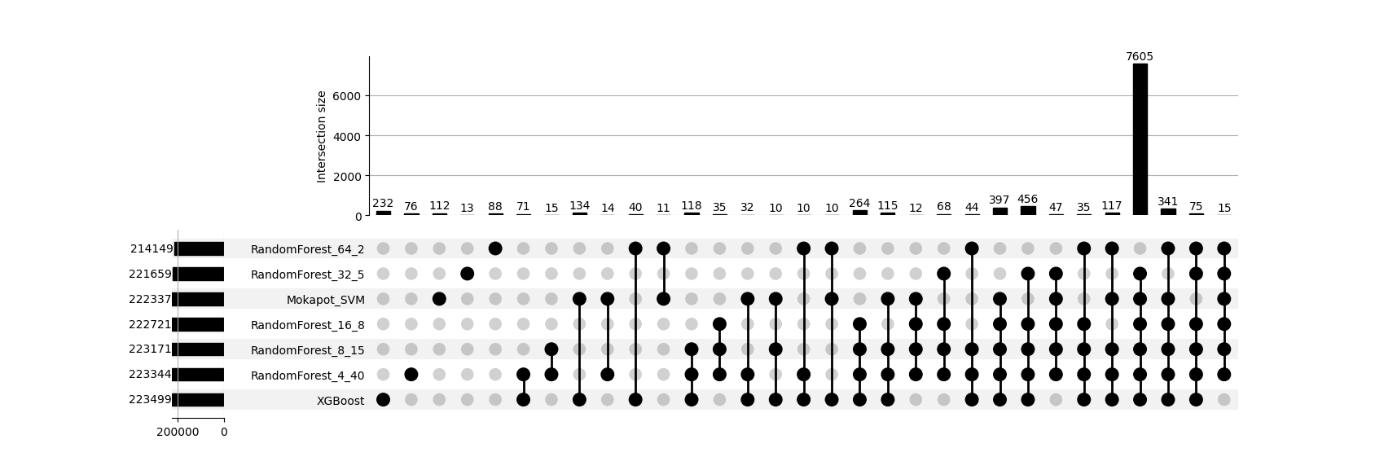

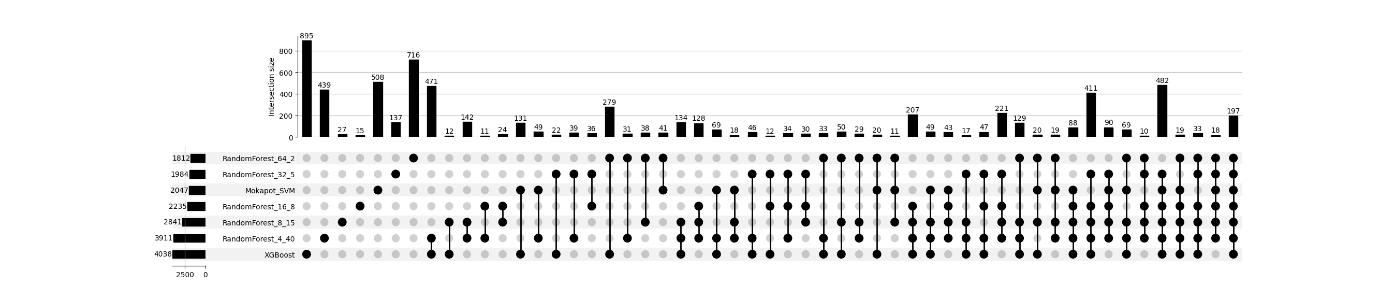

PXD004948

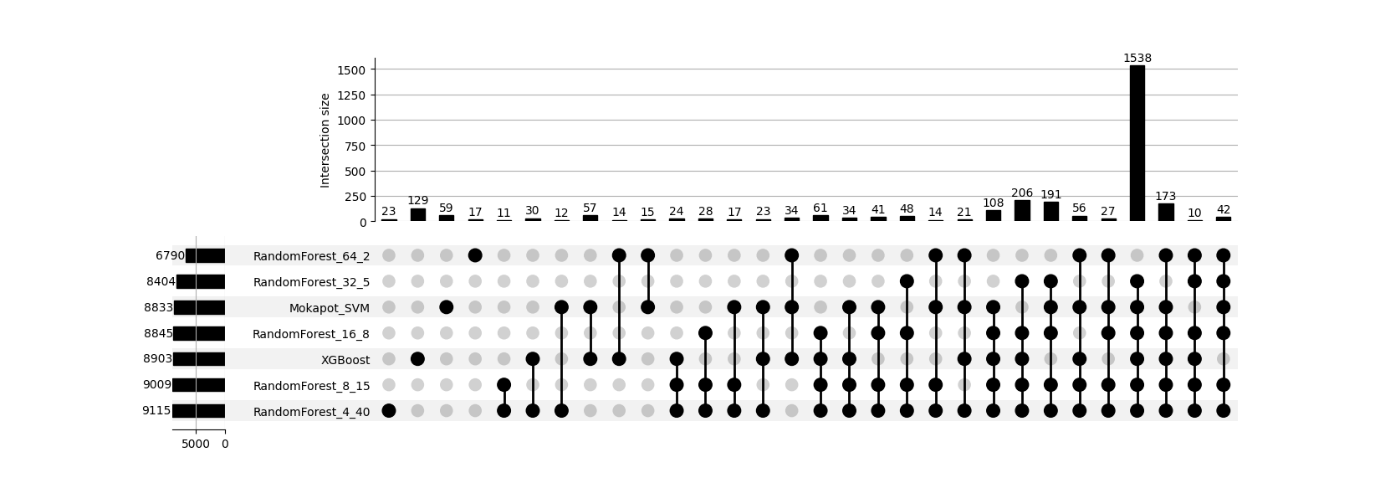

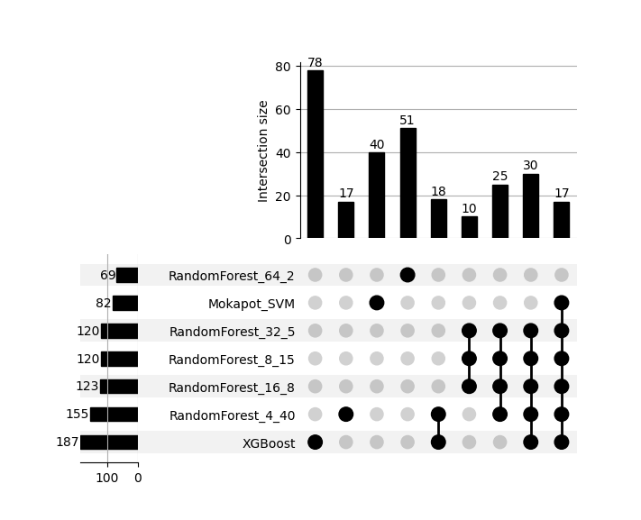

Supplementary Figure 23: UpSet plots illustrating the intersection of identified PSMs across different rescoring algorithms for MS-GF+ search results across datasets PXD000612, and PXD004948. The top panel represents target PSMs, while the down panel shows entrapment PSMs. The x-axis of each plot displays unique and shared intersections among the search engines, while the y-axis indicates the intersection size. The results highlight a strong overlap among target PSM identifications across search engines, whereas entrapment PSMs appear mostly unique to each search engine, suggesting that most entrapment matches are randomly distributed. For the target PSMs figure, the intersection of all search engines, containing the largest number of PSMs, was excluded to improve visualization and facilitate comparison of smaller intersections.

PXD040344

PXD004947

Supplementary Figure 24: UpSet plots illustrating the intersection of identified PSMs across different rescoring algorithms for MS-GF+ search results across datasets PXD040344, and PXD004947. The top panel represents target PSMs, while the down panel shows entrapment PSMs. The x-axis of each plot displays unique and shared intersections among the search engines, while the y-axis indicates the intersection size. The results highlight a strong overlap among target PSM identifications across search engines, whereas entrapment PSMs appear mostly unique to each search engine, suggesting that most entrapment matches are randomly distributed. For the target PSMs figure, the intersection of all search engines, containing the largest number of PSMs, was excluded to improve visualization and facilitate comparison of smaller intersections.

PXD004565

PXD004325

Supplementary Figure 25: UpSet plots illustrating the intersection of identified PSMs across different rescoring algorithms for MS-GF+ search results across datasets PXD004565, and PXD004325. The top panel represents target PSMs, while the down panel shows entrapment PSMs. The x-axis of each plot displays unique and shared intersections among the search engines, while the y-axis indicates the intersection size. The results highlight a strong overlap among target PSM identifications across search engines, whereas entrapment PSMs appear mostly unique to each search engine, suggesting that most entrapment matches are randomly distributed. For the target PSMs figure, the intersection of all search engines, containing the largest number of PSMs, was excluded to improve visualization and facilitate comparison of smaller intersections.

PXD009815

Supplementary Figure 26: UpSet plots illustrating the intersection of identified PSMs across different rescoring algorithms for MS-GF+ search results across dataset PXD009815. The top panel represents target PSMs, while the down panel shows entrapment PSMs. The x-axis of each plot displays unique and shared intersections among the search engines, while the y-axis indicates the intersection size. The results highlight a strong overlap among target PSM identifications across search engines, whereas entrapment PSMs appear mostly unique to each search engine, suggesting that most entrapment matches are randomly distributed. For the target PSMs figure, the intersection of all search engines, containing the largest number of PSMs, was excluded to improve visualization and facilitate comparison of smaller intersections.
